# On the flexibility of the cellular amination network in *E. coli*

**DOI:** 10.1101/2022.01.25.477661

**Authors:** Helena Schulz-Mirbach, Alexandra Müller, Tong Wu, Pascal Pfister, Selçuk Aslan, Lennart Schada von Borzyskowski, Tobias J. Erb, Arren Bar-Even, Steffen N. Lindner

## Abstract

Ammonium (NH4^+^) is essential to generate the nitrogenous building blocks of life. It gets assimilated via the canonical biosynthetic routes to glutamate and is further distributed throughout metabolism via a network of transaminases. To study the flexibility of this network, we constructed an *Escherichia coli* glutamate auxotrophic strain. This strain allowed us to systematically study which amino acids serve as amine source and found that several amino acids complement the auxotrophy, either by producing glutamate via transamination reactions or by their conversion to glutamate. In this network, we identified aspartate transaminase AspC as a major connector between many amino acids and glutamate. Additionally, we extended the transaminase network by the amino acids β-alanine, alanine, glycine and serine as new amine sources and identified D-amino acid dehydrogenase (DadA) as an intracellular amino acid sink removing substrates from transaminase reactions. Finally, ammonium assimilation routes producing aspartate or leucine were introduced. Our study reveals the high flexibility of the cellular amination network, both in terms of transaminase promiscuity and adaptability to new connections and ammonium entry points.

## Introduction

Nitrogen is essential for all forms of life, as it is part of 75 % of the cells building blocks (mainly in proteins and nucleic acids) (Milo & Phillips, 2015). The conversion of atmospheric dinitrogen (N_2_) to ammonia (NH_3_) by diazotrophic bacteria or industrially by the Haber–Bosch process is essential to make it available for the assimilation by plants and other organisms to produce nitrogenous compounds.

While carbon fixation has evolved several times, resulting in versatile naturally occurring ways of carbon fixation (Löwe & Kremling, 2021), the introduction of ammonium (NH_4_^+^, protonated form of ammonia) into the building blocks of life is similar in all organisms and limited to the fixation of ammonium at the node between 2-oxoglutarate, glutamate and glutamine (Fig. 1). Here, three cooperating canonical enzymes assimilate ammonium in two distinct ways. In the first reaction, glutamate is the direct product of glutamate dehydrogenase (*gdhA*, GDH). In the second pathway, the combined activity of glutamine synthetase (*glnA*, GS) and glutamate synthase (Glutamine 2-oxoglutarate aminotransferase, *gltBD*, GOGAT) fix another ammonium and convert glutamate into glutamine, which then donates one amine to 2-oxoglutarate to form two glutamate molecules (Helling, 1994; Kumar & Shimizu, 2010) (Fig. 1). In order to make all essential amino acids and other aminated compounds, glutamate and glutamine then donate their amines to specific keto acids or other amino acid precursors in reversible transferase reactions. As ammonium enters metabolism solely via these two routes, all cellular nitrogen is provided by either glutamate (75%) or glutamine (25%) (Yang et al, 2018).

**Figure 1:**
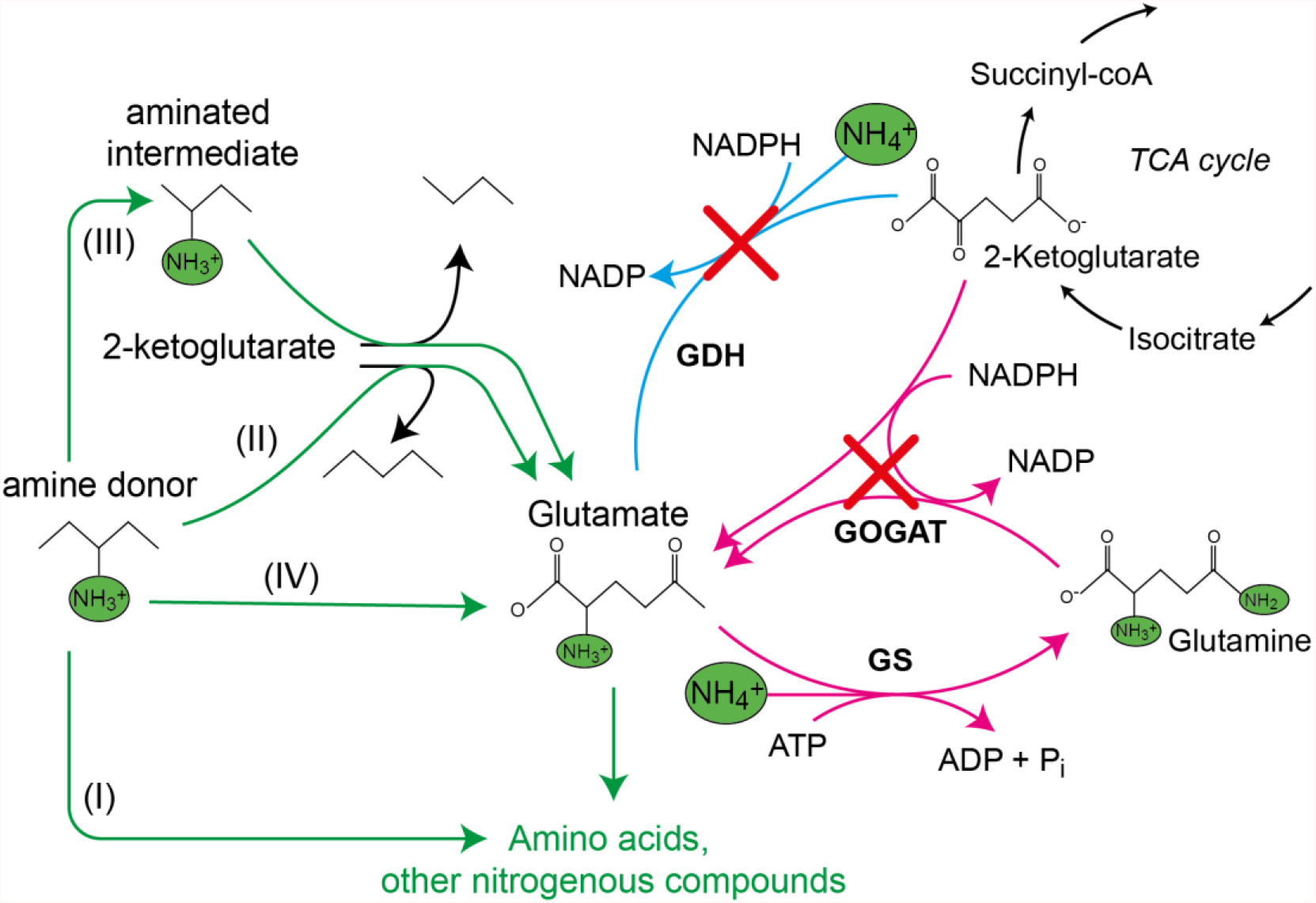
Canonical ammonium assimilation via glutamate dehydrogenase (GDH) (blue arrow) or glutamine synthetase (GS) and glutamine 2-oxoglutarate aminotransferase (GOGAT) (pink arrows). One pathway for ammonium assimilation is the amination of the TCA cycle intermediate 2-ketoglutarate by GDH to form glutamate (blue arrow). A second pathway requires joint action of GS and GOGAT, which first aminate glutamate to form glutamine (GS), which donates one amine to 2-ketoglutarate (GOGAT) to form two glutamate molecules, which further provide amines (green) for biosynthesis of amino acids and other nitrogenous compounds (pink arrows). Growth of the glut-aux strain deleted in GDH and GOGAT (red crosses) by a supplemented amine source is possible via the following mechanism. The amine donor either (I) replaces glutamate as amine source for production of amino acids and nitrogenous compounds, (II) donates an amine to 2-ketoglutarate to form glutamate, (III) is converted to an intermediate donating an amine to 2-ketoglutarate or (IV) is metabolically converted into glutamate. Green arrows indicate these cases.

Besides the glutamate biosynthesis node, alternative entry points for ammonium theoretically exist, e.g. alanine dehydrogenase or aspartate ammonia lyase, but these are not relevant for ammonium assimilation (Kim & Hollocher, 1982). Evolution has developed a system for ammonium assimilation which is controlled by its intracellular availability. The ATP investment driving GS activity makes amination reactions favorable even at low ammonium concentrations. Additionally, most of the GS orthologues have evolved kinetic parameters optimized for lower ammonium concentrations with an apparent *K*_*M*_ of 0.1 mM for ammonium (Reitzer, 2014). At high ammonium concentrations, the NADPH dependent (and thus energetically cheaper) GDH allows more efficient ammonium assimilation than GS (Reitzer, 2014). This metabolic switch might explain the prominence of glutamate-based ammonium assimilation as opposed to other ammonia entry points in nature. To generate the NADPH required for GDH, *Escherichia coli* mostly uses the membrane-bound proton-translocating transhydrogenase (PntAB) (Sauer et al, 2004). This enzyme exploits the proton motif force to drive proton translocation from NADH to NADP^+^ (Spaans et al, 2015), and thereby indirectly competes with ATP synthesis. Therefore, when growing under high ammonium concentrations, growth of this microorganism might benefit from ammonium assimilation via NADH dependent dehydrogenases.

Following these thoughts, we tried to assess if alternative routes for ammonium assimilation can arise from the metabolic network of *E. coli*. For this purpose, we systematically investigated the flexibility of the amination network in a glutamate auxotrophic -and hence ammonium assimilation deficient-*E. coli* strain. This study provides fundamental knowledge on the plasticity of ammonium metabolism in *E. coli* and moreover addresses industrial interests by providing a versatile bacterial *chassis* for screening and optimization of ammonium assimilation and transamination reactions.

## Results

### Only some amino acids serve as amine source

To study the flexibility of *Escherichia coli*’s cellular amination network, we first generated a strain in which both canonical ammonia assimilation routes are disrupted. Accordingly, we deleted the genes encoding GDH (*gdhA*) and GOGAT (*gltBD*), which are responsible for 2-oxoglutarate amination under high and low ammonia concentrations, respectively (Helling, 1994; Kumar & Shimizu, 2010) (Fig. 1). The resulting glutamate auxotrophic strain (glut-aux, Δ*gdhA* Δ*gltBD*) was not able to grow in minimal medium with ammonia as sole nitrogen source unless an amine group donor like glutamate was provided in the medium (Fig. 2). Initially, we were interested in testing whether other amino acids can replace glutamate as an amine source allowing growth of the glut-aux. We therefore characterized growth of the glut-aux strain when supplemented with one of the proteinogenic or naturally occurring non-proteinogenic amino acids (ornithine and β-alanine). We note that this growth experiment is different from the experiments commonly described in literature, where amino acids were added to the medium without ammonia to serve as sole nitrogen source (Neidhardt et al, 1996). In these experiments, the metabolic degradation of the amino acids to release ammonia suffices to enable growth. Conversely, for the glut-aux strain, growth complementation through the supplemented amino acids as amine source must follow one of these options: (i) replace glutamate as an amine donor for the production of other nitrogenous compounds; (ii) donate their amine group to 2-oxoglutarate to generate glutamate as an amine; (iii) be metabolically converted to compounds that can donate their amine group to 2-oxoglutarate; or (iv) be metabolically converted to glutamate (Fig. 1). In the former three cases, the amination network of the cell needs to be flexible enough to adapt to different directionalities of at least some of the transamination reactions. As utilization of amino acids as amine group donors in the glut-aux strain might be dependent on the nitrogen regulated (Ntr) response (Reitzer, 2003), we performed growth experiments with and without ammonia in the medium (light blue and magenta lines in Fig. 2). As a control, we repeated the classical experiments of testing each amino acid as sole nitrogen source with a wild type strain. Here, amino acids are not required to directly donate their amine group but can rather support growth by releasing ammonia through their degradation (black lines in Fig. 2).

**Figure 2:**
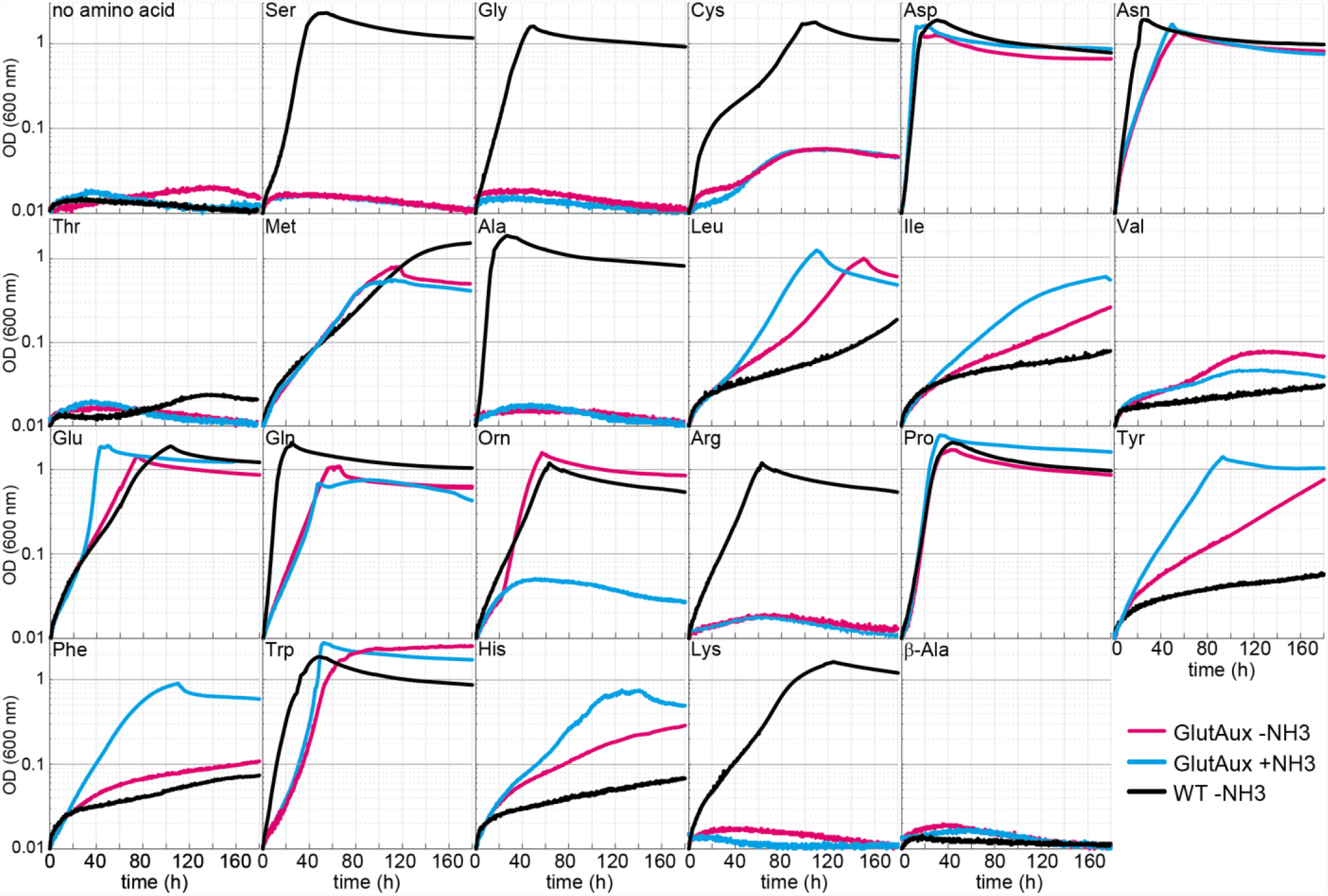
Identification of amino acids which rescue growth of the glut-aux strain. The glut-aux strain was grown in M9 medium with (blue line) or without ammonium (magenta line) and 20 mM glycerol as carbon source. *E. coli* WT was grown in M9 medium without ammonium (black line) and 20 mM glycerol as carbon source. 5 mM of the indicated amino acids or no amino acid as negative control were supplemented to test if they can serve as an amine source (glut-aux strain) or an ammonium source (WT). Data shows representative growth as observed from triplicate repeats with errors < 5 %.

We found that only some amino acids rescue growth of the glut-aux strain (Fig. 2). This generally correlated with the existence of known transaminase enzymes that enable glutamate production from the respective amino acids in *E. coli*. For example, aspartate, leucine, and tyrosine serving as cellular amine donor for glutamate generation from 2-oxoglutarate could be attributed to the activity of aspartate transaminase (AspC), tyrosine transaminase (TryB), and branched-chain-amino-acid transaminase (IlvE) (Gelfand & Steinberg, 1977). As these transaminases display considerable cross reactivity (Gelfand & Steinberg, 1977; Inoue et al, 1988), each of these three amino acids might support the production of the others directly, without the need for glutamate as an amine donor. However, glutamate here still likely serves as the primary amine donor for most cellular nitrogenous compounds (Yang et al, 2018). Hence, AspC, TryB, and IlvE must also be fully reversible under physiological conditions to aminate 2-oxoglutarate to glutamate. While transaminases are generally reversible enzymes, their ability to effectively operate reversibly *in vivo* is not trivial as the [glutamate]/[2-oxoglutarate] ratio is very high (above 100) under physiological conditions (Bennett et al, 2009), making the reverse amine transfer onto 2-oxoglutarate to form glutamate thermodynamically challenging. Since the glut-aux strain grew with several amino acids as amine source, we conclude that the cellular amination network must be sufficiently flexible to accept amine sources other than glutamate despite the potential thermodynamic barriers.

To prove amine transfer from the provided amino acids, we cultivated the glut-aux strain with 5 mM of one of five (unlabeled) representative amino acids that can serve as an amine donor – glutamate, proline, aspartate, tryptophan, and leucine – in a medium containing 20 mM ^15^N-ammonium. We subsequently measured the ^15^N labeling in proteinogenic amino acids (Methods) and found that most of them were completely unlabeled (Fig. 3A), confirming that their amine group was transferred from the amino acid rather than from free (^15^N-labeled) ammonia in the medium (Fig. 3B). Arginine was once labeled (R, Fig. 3), as one of its nitrogen atoms originates from the amide nitrogen of glutamine that is derived from ammonia fixed by GS activity (Fig. 3B). Overall, these results confirm that the amino acids added to the medium were the only amine sources allowing growth of the glut-aux strain, rather than allowing amination of amino acid derived backbones with free ammonium in the medium.

**Figure 3:**
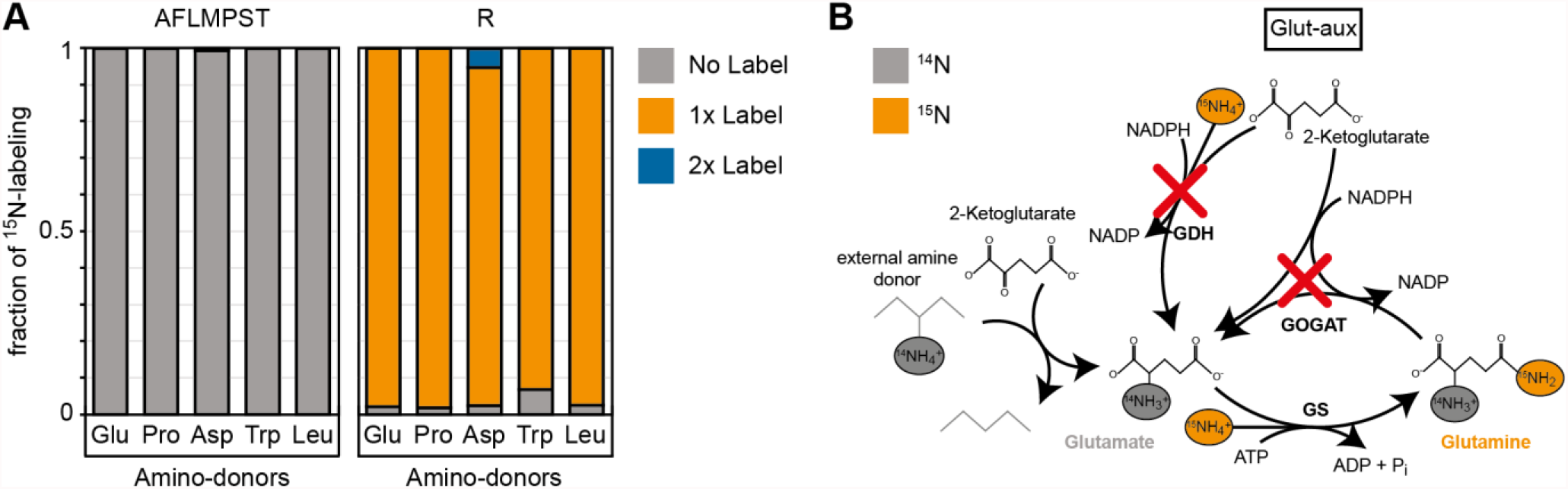
^15^N-labeling confirms amine assimilation from supplied amino acid in the glut-aux strain. **A** The glut-aux strain was incubated in ^15^N-NH_3_Cl M9 medium with 20 mM glycerol as carbon source. ^15^N labeling pattern in analyzed proteinogenic amino acids (single letter code) upon feeding with 5 mM of unlabeled amino acids glutamate, proline, aspartate, tryptophan, or leucine as amine-donors (three letter code). The labeling pattern of the amino acids A, F, L, M, P, S and T were identical with supplied amino acids and hence only a representative dataset is shown. Data represents means of triplicate measurements with errors < 5 %. **B** Schematic presentation of expected ammonium incorporation in glut-aux grown on M9 with ^15^N-NH_3_Cl with 20 mM glycerol as carbon source and an unlabeled amino acid as ammonium source. Since the genes encoding GDH and GOGAT are deleted in the glut-aux (red crosses), the glut-aux thus relies on the provided amino acid for biosynthesis of unlabeled (grey) glutamate. During glutamine biosynthesis, the glut-aux assimilates ^15^N-NH_3_Cl to form once labelled glutamine and thus once labelled arginine.

To further validate that growth of the glut-aux strain is limited by the supply of amine groups through the amino acids provided in the medium we cultivated it using different concentrations of amino acids (Fig. 4). As expected, we found that biomass yield, as indicated by the maximal OD_600_, directly correlated with the concentration of the supplemented amino acid. All amino acids showed the same dependency of biomass yield on concentration, with the exception of ornithine, which supported roughly double the yield for each concentration. This is in line with the fact that both amine groups from ornithine can be donated to aminate 2-oxoglutarate to glutamate (Prieto-Santos et al, 1986). As this ornithine degradation pathway is induced by nitrogen starvation (Schneider et al, 2013; Schneider & Reitzer, 2012), growth of the glut-aux strain with ornithine as amine group donor was observed only when ammonia was omitted in the medium. The correlation of maximal OD_600_ to amino acid concentration confirms that amine supply from the amino acid limits biomass yield in the glut-aux strain in the same manner for all tested amino acids. To our surprise glutamate was not the amine donor supporting fastest growth of the glut-aux strain. Even proline, which in order to donate its amine group needs to be converted to glutamate, supported faster growth (Fig. 4). This, together with the fact that the growth rate increased proportionally with the glutamate concentration indicates that glutamate uptake is limiting growth of the glut-aux strain.

**Figure 4:**
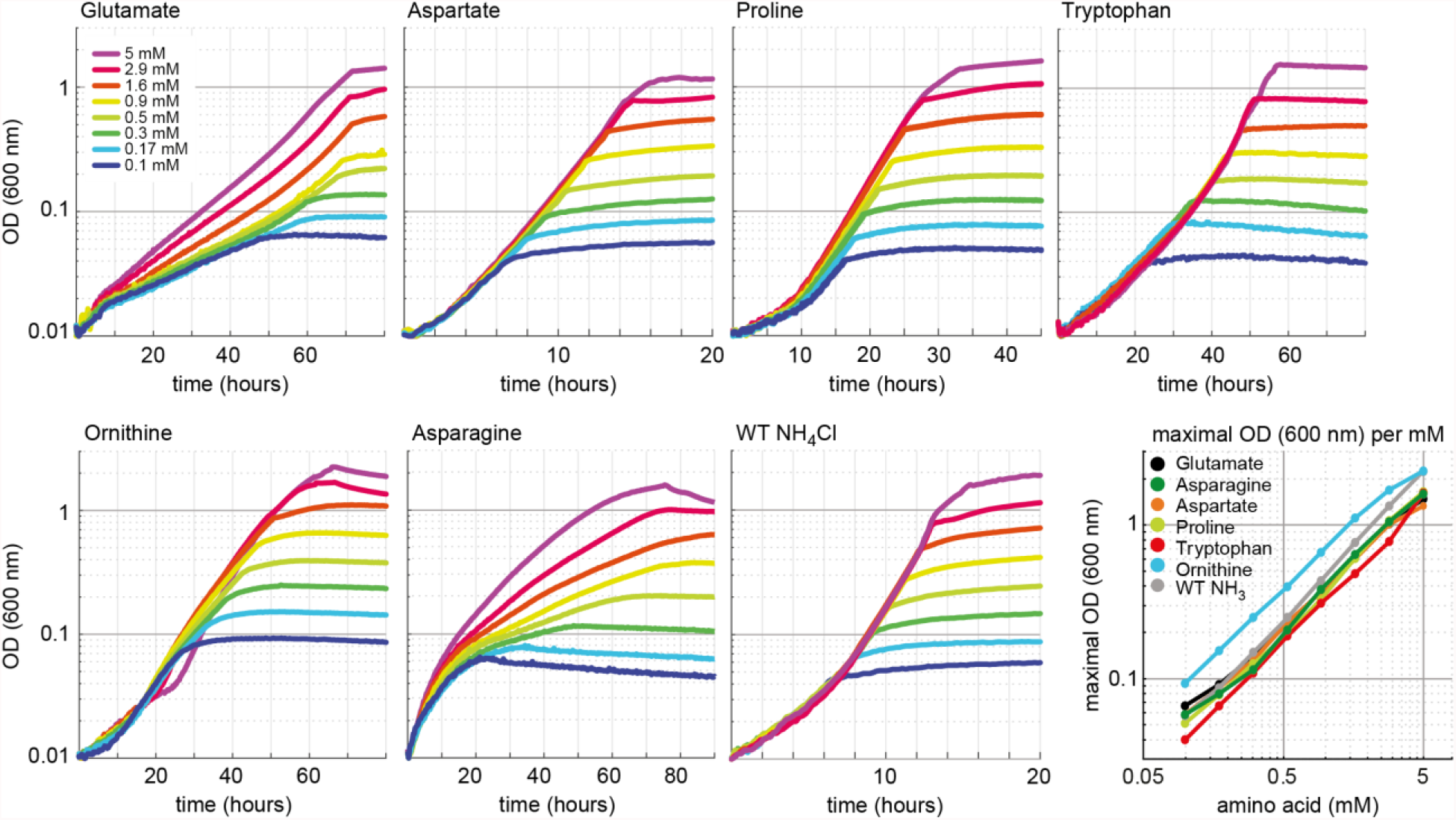
Growth dependency of the glut-aux strain on amino acid concentration. Cells were grown in ammonium free M9 medium with 20 mM glycerol and the indicated concentrations of the amino acids glutamate, aspartate, proline, tryptophan, ornithine and asparagine. As a comparison WT was grown in ammonium free M9 medium with 20 mM glycerol but with NH_4_Cl concentrations similar to the amino acid concentrations used. Data shows representative growth as observed from triplicate repeats with errors < 5 %.

### The cellular amination network is highly promiscuous

To investigate the contribution of different enzymes to the use of amino acids as amine source, we decided to analyze effects of several gene deletions. We specifically deleted genes encoding proteins essential for the certain degradation pathways allowing glutamate formation based on available documentation in the database Ecocyc (Keseler et al, 2021). First, we explored use of proline as amine source (Fig. 2). We did not expect this amino acid to donate its amine group directly, but rather to be metabolized by PutA, encoding for a bifunctional flavoenzyme with proline dehydrogenase and 1-pyrroline-5-carboxylate dehydrogenase activities (Moxley et al, 2014), to glutamate (Frank & Ranhand, 1964) which would serve as an amine donor. Indeed, upon deletion of *putA* growth of the WT and of the glut-aux strain on proline as sole ammonium source was abolished (Fig. 5A). This confirmed that proline could only support growth via its native degradation pathway and was unable to serve as amine source via a different pathway. We then focused on amino acids that are expected to directly donate their amine groups. For example, methionine, which could serve as an amine donor, was previously found to be the main substrate for only a single transaminase: YbdL (Dolzan et al, 2004). However, deletion of *ybdL* in the Δ*gdhA* Δ*gltBD* strain did not affect growth with methionine as amine donor (Supplementary Fig. S1), suggesting that the physiological contribution of this transaminase to the use of methionine as amine source is negligible. YbdL was also shown to be the only transaminase able to efficiently accept histidine and phenylalanine (Dolzan et al, 2004; Inoue et al, 1988). Yet, the glut-aux Δ*ybdL* strain did not show any growth retardation when using histidine or phenylalanine as amine donor (Supplementary Fig. S1). Indeed, other transaminases are known to accept methionine and histidine, albeit at a substantially lower affinity and rate than their primary substrate, e.g., aspartate transaminase, tyrosine transaminase, and branched-chain-amino-acid transaminase (Inoue et al, 1988; Mavrides, 1987; Powell & Morrison, 1978). It therefore seems that the promiscuity of such transaminases enables effective use of methionine as amine donor even in absence of the enzyme preferring it as substrate.

**Figure 5.**
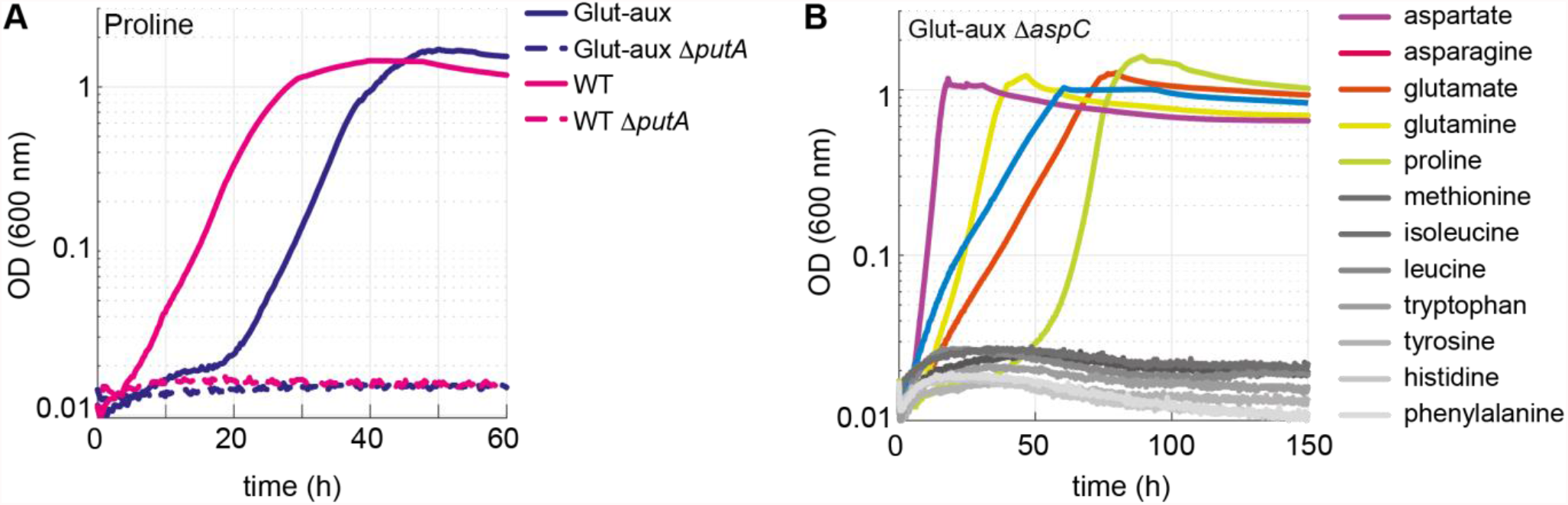
Proline is converted to glutamate to serve as an amine source. AspC is responsible for the utilization of several amino acids. **A** Deletion of *putA* abolished growth with proline as an amine source in the glut-aux strain and as a nitrogen source in the WT. **B** Deletion of *aspC* eliminates growth of the glut-aux with methionine, leucine, isoleucine, histidine, tyrosine, tryptophan and phenylalanine as amine donor (grey lines). Experiments were carried out in M9 w/ammonium containing 20 mM glycerol and 5 mM of the indicated amine sources. Data represents means of triplicates with < 5 % variation.

To further investigate the relevance of promiscuous transaminase activities for using external amine donors, we focused on AspC. This transaminase is known to accept a range of amino acids. After constructing the glut-aux Δ*aspC* strain, its growth was analyzed on all previously tested 22 amino acids. We found the glut-aux Δ*aspC* strain to be unable to grow with histidine, tyrosine, phenylalanine, tryptophan, methionine, isoleucine and leucine as amine donors, which all allowed growth of the glut-aux strain (grey lines in Fig. 5B, Fig. 2). This demonstrates promiscuity of AspC and reveals that, in its physiological context, this transaminase is even more versatile than previously reported. Although AspC had shown low specific transaminase activity with histidine, methionine, isoleucine and leucine *in vitro* (Hayashi et al, 1993), our experiments suggest, however, that the enzyme is essential for utilization of these amino acids as amine donors *in vivo*. Furthermore, TyrB and IlvE, which had shown higher specific *in vitro* activities with tyrosine, phenylalanine and tryptophan compared to AspC (Hayashi et al, 1993), apparently did not complement the AspC deletion.

### Engineering utilization of alanine as amine donor

Despite the presence of two glutamate producing alanine transaminases (AlaA, AlaC) as well as multiple transaminases that can promiscuously accept alanine (Kim et al, 2010; Pena-Soler et al, 2014), alanine did not serve as amine donor for the glut-aux strain, regardless of the presence or absence of ammonia in the medium (Fig. 2). Even upon overexpression of the alanine transaminase isozymes AlaA and AlaC, no growth with alanine as amine donor was observed (Fig. 6A). Thus, we employed adaptive laboratory evolution to enable the glut-aux strain to use alanine as an amine donor. We incubated the glut-aux strain with and without overexpression of either AlaA or AlaC on 20 mM glycerol + 5 mM alanine. After two weeks of incubation the glut-aux cultures overexpressing AlaA had grown to an OD_600_ > 1, while the cultures which did not overexpress any alanine transaminases had not grown at all. The grown cultures were used to isolate single colonies, which upon transfer to M9 medium with 20 mM glycerol and 5 mM alanine grew immediately, indicating that indeed mutations allowed their growth (Fig. 6A). To analyze which genetic differences are present in the evolved strains, we sent two of the independently obtained isolates and the parent strain for genome sequencing. Analysis of the genome sequencing results revealed that both isolates had mutations in the alanine racemase gene *dadX* (Supplementary Table S1, Supplementary Figure S2), either a duplication of a 27 bp region from nucleotide 151 to nucleotide 177 (mutant 1) or a 1 bp deletion at nucleotide position 821 (of the 1071 nucleotide gene) (mutant 2). In both cases the mutations resulted in a frameshift, indicating a loss of function of DadX. Alanine racemase diverts flux from L-alanine to D-alanine which is either used for cell wall biosynthesis (Walsh, 1989) or degraded to pyruvate via D-alanine:quinone oxidoreductase (DadA) (Franklin & Venables, 1976). The absence of DadX activity likely reduces flux into an important alanine sink, resulting in higher alanine availability which might be responsible for growth rescue in the mutants. To verify this hypothesis, we deleted *dadX* in the evolved strain, as well as in the parental glut-aux strain. Both strains (with overexpression of AlaA) immediately grew with alanine as amine donor (Fig. 6A, Supplementary Figure S3), verifying that reducing an important intracellular alanine sink allows the overexpressed transaminase to use alanine as an amine donor to effectively sustain growth. Noteworthy, a deletion of DadA in the glut-aux strain overexpressing AlaA did not restore growth with alanine as amine donor, indicating that the reversible conversion of L-alanine to D-alanine is already sufficient to reduce alanine availability below a critical level, where it cannot support enough flux via AlaA for cellular growth. Note that *dadX* is not essential, as *E. coli* possesses additional genes that are able to provide sufficient D-alanine for cell wall biosynthesis (Kang et al, 2011; Wild et al, 1985).

**Figure 6.**
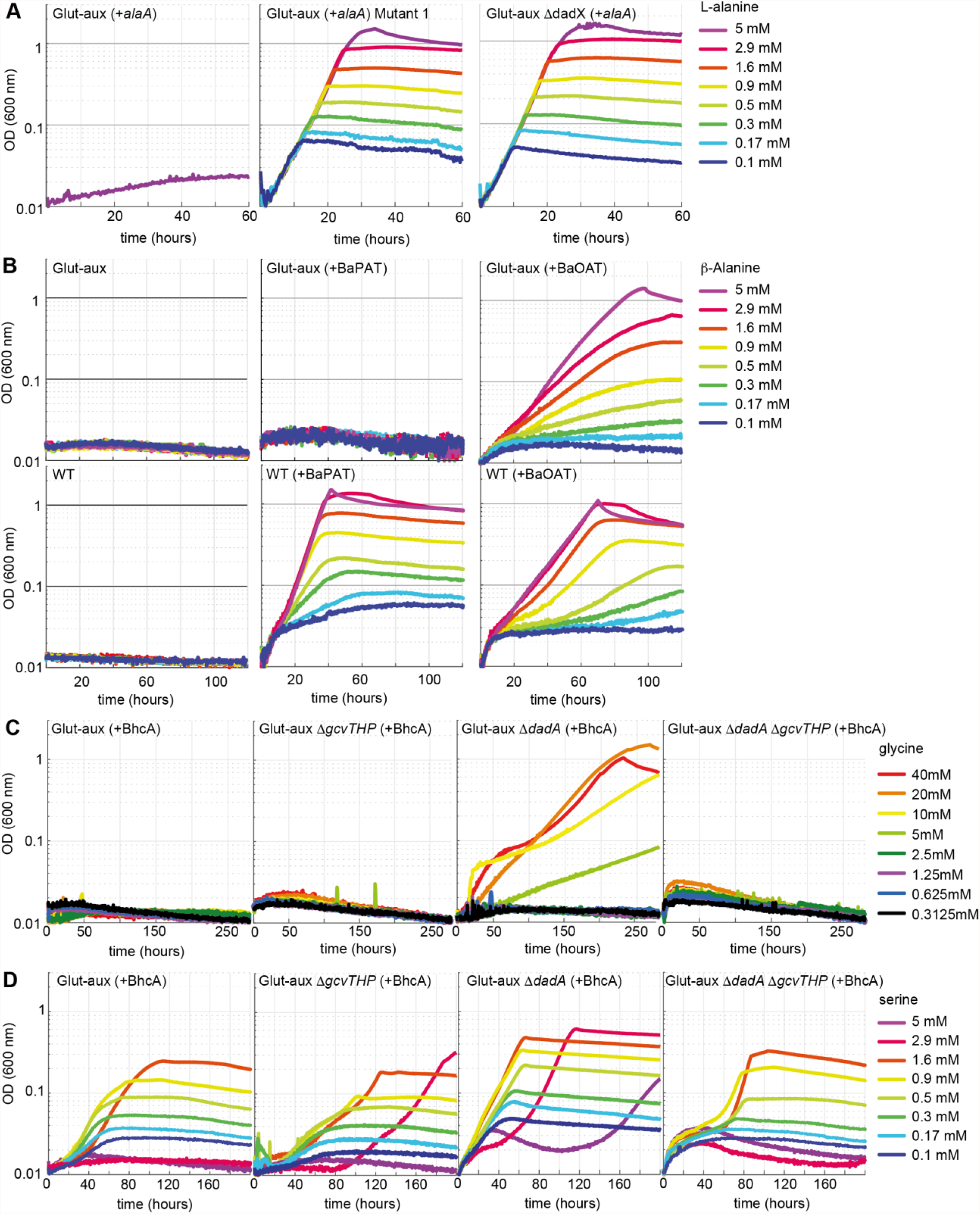
Glut-aux strains engineered to use alanine, β-alanine, glycine and serine as amine source. **A** DadX mutation or deletion and alanine transaminase overexpression allows alanine utilization as amine source. Growth of the glut-aux strains on 20 mM glycerol with 5 mM alanine as amine donor. **B** Overexpression of β-alanine-2-oxoglutarate transaminase (BaOAT) enables amine donation from β-alanine in the glut-aux strain. Glut-aux strain and WT overexpressing β-alanine-pyruvate aminotransferase (BaPAT) or β-alanine-2-oxoglutarate aminotransferase (BaOAT) where grown in media containing no N-source (WT) or ammonium-chloride (glut-aux), 20 mM glycerol and the indicated β-alanine concentrations. **C** DadA deletion and BhcA overexpression allows glycine utilization as amine donor. Growth of the glut-aux strains +BhcA on 20 mM glycerol with indicated glycine concentrations. Data represents triplicate measurements with < 5 % variation. **D** BhcA overexpression allows use of serine as amine source, which is improved by additional deletion of *dadA*. Growth of the glut-aux strains +BhcA on 20 mM glycerol with indicated serine concentrations. All data represents triplicate measurements with < 5 % variation.

### Engineering the utilization of β-alanine as amine donor

The non-proteinogenic amino acid β-alanine is neither a suitable amine donor for the glut-aux strain nor an N-source for the wildtype (Fig. 2). In order to test if we could engineer the glut-aux strain to use β-alanine as an amine donor, we tested overexpression of two different β-alanine transaminases, the β-alanine-2-oxoglutarate aminotransferase from *Saccharomyces kluyveri* (BaOAT, Uniprot ID A5H0J5) and the β-alanine-pyruvate aminotransferase from *Pseudomonas aeruginosa* (BaPAT, Uniprot ID Q9I700). Overexpression of either BaPAT or BaOAT allowed the wildtype strain to grow using β-alanine as sole source of ammonia (Fig. 6B). The enzymes transfer the amine group from β-alanine onto the ketoacids pyruvate and 2-oxoglutarate to generate alanine and glutamate, respectively. Both alanine and glutamate serve as N-sources for the wildtype (Fig. 2). The glut-aux strain however was only able to grow with β-alanine as an amine donor, when BaOAT, but not when BaPAT was overexpressed (Fig. 6B), which is in good agreement with our previous results that glutamate, but not alanine, can immediately serve as an amine donor in the glut-aux strain (Fig. 2). To further confirm that growth of the glut-aux + BaOAT strain was rescued by directly using β-alanine via the proposed transamination reaction catalyzed by BaOAT, we conducted a nitrogen-tracing experiment. The medium contained ^15^N-labeled ammonium together with 20 mM glycerol and 5 mM β-alanine. As expected, the majority of amino acids did not show any ^15^N-labeling (Supplementary Figure S4), confirming that their amino group originated from β-alanine, rather than from ammonium. The only exception were arginine and glutamine that were single-labeled, with the ^15^N-label originating from glutamine synthetase activity (see above).

### Overexpression of a glycine-oxaloacetate transaminase allows growth of the glut-aux strain with glycine after deletion of a glycine sink

Like alanine, glycine did not serve as amine source in the glut-aux strain (Fig. 2). As *E. coli* lacks a glycine transaminase, this amino acid cannot be directly generated by transamination of the respective ketoacid (glyoxylate). Instead, it is obtained from serine via the serine-hydroxymethyl-transferase reactions. To engineer the usage of glycine as amine donor, we overexpressed the glycine-oxaloacetate transaminase from *Paracoccus denitrificans* (BhcA, Uniprot ID A1B8Z3) in the glut-aux strain to form aspartate, which we have shown to support fast growth of the glut-aux strain (Fig. 2).

However, growth of the glut-aux strain overexpressing BhcA was not restored upon supplying the medium with glycine (Fig. 6 C, Supplementary Figure S5). We speculated that, similar to our results with alanine as amine donor described above, the provided glycine is further diverted into a sink. Hence we directed our efforts towards the glycine cleavage system and the potential promiscuous activity of the D-alanine:quinone oxidoreductase DadA with glycine. DadA has been reported to be active with a variety of D-amino acids (Wild & Klopotowski, 1981), and might potentially use glycine as a substrate as well. Indeed, the deletion of *dadA* allowed the strain to grow slowly with glycine as an amine donor (Fig. 6C), at first indicating glycine removal via promiscuous activity of DadA. However, *in vitro*, DadA showed only a very low specific activity with glycine (Supplementary Figure S6A) and an apparent *K*_M_ > 500 mM for this amino acid (Supplementary Table S2), which strongly suggested that the catalytic efficiency (*k*_cat_/*K*_M_ of 0.275 M^-1^ s^-1^) of the enzyme (Supplementary Figure S6A, Supplementary Table S2) was insufficient to provide a strong glycine sink *in vivo*. Notably, DadA was previously reported to efficiently convert D-serine (Wild & Klopotowski, 1981), the product of alanine racemase Alr, which also acts on L-serine besides L-alanine (Ju et al, 2005). *In vitro* measurements of DadA confirmed a higher catalytic efficiency (*k*_cat_/*K*_M_ of 5.71 M^-1^ s^-1^) of DadA with with D-serine (Supplementary Figure S6A) than previously measured with glycine. BhcA exhibits much better kinetic properties with serine (tested with glyoxylate as the acceptor (*K*_M_ 2.1 mM; *k*_cat_ 8.8 s^-1^)) than with glycine (Schada von Borzyskowski et al, 2019). We thus concluded that glycine needed to be converted to serine via combined activities of the glycine cleavage system (GCV), cleaving one glycine molecule to ammonium, CO_2_, and 5,10-methylene-tetrahydrofolate, and serine-hydroxymethyl-transferase subsequently condensing the latter with a second glycine molecule to form serine. This pathway is used by *E. coli* when utilizing glycine as sole source of nitrogen (Newman et al, 1976). Supporting this hypothesis, the deletion of the GCV subunits *gcvTHP* in the glut-aux strain overexpressing BhcA did not allow growth with glycine. Moreover, deletion of *gcvTHP* in the glut-aux Δ*dadA* strain overexpressing BhcA abolished growth with glycine (Fig. 6C), providing strong evidence that glycine is converted to serine to utilize it as amine donor (Supplementary Figure S7). Moreover, compared to other amine donors tested the growth supported by glycine was very slow (stationary phase reached after 200 h).

Next, we tested if serine (not an amine donor in the glut-aux strain (Fig 2)), served as amine donor for the glut-aux strain overexpressing BhcA. In good agreement with the conclusion of our glycine trials we found that growth of the glut-aux, glut-aux Δ*dadA*, glut-aux Δ*gcvTHP* and the glut-aux Δ*dadA* Δ*gcvTHP* strains (all overexpressing BhcA) was restored upon addition of low serine concentrations (Fig. 6D). In our growth experiments, serine concentrations above 1.7 mM seemed to be toxic to the strain, which can be explained by inhibition of isoleucine and aromatic amino acid biosynthesis by serine derived hydroxypyruvate (Hama et al, 1990). The glut-aux Δ*dadA* reached the highest OD_600_ on all tested serine concentrations, indicating that also here deletion of DadA is beneficial as it removes a sink for either glycine or serine directly (Fig. 6D). To further demonstrate serine transamination catalyzed by BhcA with oxaloacetate as amine acceptor, we conducted *in vitro* measurements with purified BhcA. The enzymatic coupling assay indeed confirmed amine transfer from serine to oxaloacetate (Supplementary Figure S6B, Supplementary Table S3). The resulting *K*_M_ of 7 mM and a *k*_cat_ of 25.9 s^-1^ (Supplementary Table S3) indicated a sufficient turnover of serine and oxaloacetate into aspartate and hydroxypyruvate. To further verify that the strain is indeed using the amine from glycine or serine, respectively, we conducted a labelling experiment growing the glut-aux Δ*dadA* overexpressing BhcA and a WT control on M9 with ^15^NH_4_ or M9 with ^14^NH_4_ supplemented with 20 mM glycerol and 20 mM glycine, or 1.7 mM serine. Aspartate, proline, serine, alanine and phenylalanine were unlabeled in the glut-aux Δ*dadA* strain overexpressing BhcA, confirming that indeed all amines were derived only from the provided amino acid and not from free ammonium in the medium (Supplementary Figure S8).

### Exploring alternative routes of ammonium assimilation

After testing and engineering the glut-aux strain to use amino acids as amine group donors, we aimed to explore alternative ammonium assimilation pathways, thus a rewiring of canonical ammonium assimilation via glutamate. For this, we used the ammonium assimilation deficient glut-aux strain to test the enzymes aspartate ammonia-lyase and leucine dehydrogenase for their activity to supply all cellular amine for growth.

### Aspartate ammonia-lyase

Since aspartate served as an efficient amine donor (Fig. 2), we investigated whether it could also replace glutamate as the formation product of ammonium assimilation, catalyzed via reverse activity of aspartate ammonia-lyase (AspA; Uniprot ID P0AC38, fumarate + NH_4_ = aspartate), an *E. coli* native reaction. This enzyme canonically operates as part of the aspartate utilization/degradation pathway releasing ammonia. However, the thermodynamics of this reaction reveal a full reversibility with a Δ_r_G^’m^ of 4 [kJ/mol] (in the aspartate forming direction at a concentration of 1 mM for all reactants; http://equilibrator.weizmann.ac.il/). We hence overexpressed AspA in the glut-aux strain and analyzed its growth on a variety of carbon sources (Fig. 7A). Here, growth of the strain is dependent on both the ammonium assimilation activity to form aspartate and the subsequent aminotransferase activity to rescue the glut-aux. Growth of the glut-aux strain overexpressing AspA (+AspA) was indeed observed on all tested carbon sources (Fig. 7A), suggesting that aspartate and AspA can replace the canonical glutamate-based ammonium assimilation pathway. The glut-aux +AspA strain grew fastest on carbon sources closest to fumarate, the substrate of AspA (2-oxoglutarate, succinate, malate, fumarate). Other carbon sources (glucose, xylose, glycerol, acetate), which do not directly generate fumarate, also supported growth, but with much higher doubling times. Assimilation of free ammonium from the medium by the glut-aux +AspA strain was confirmed by a ^15^N-labelling experiment comparing proteinogenic amino acid labeling of the glut-aux +AspA strain with the WT (serving as a positive control) upon growth on 20 mM succinate with either ^15^NH_4_ or ^14^NH_4_. Both strains, the WT and the glut-aux +AspA strain, showed identical labeling (Fig. 7C).

**Figure 7:**
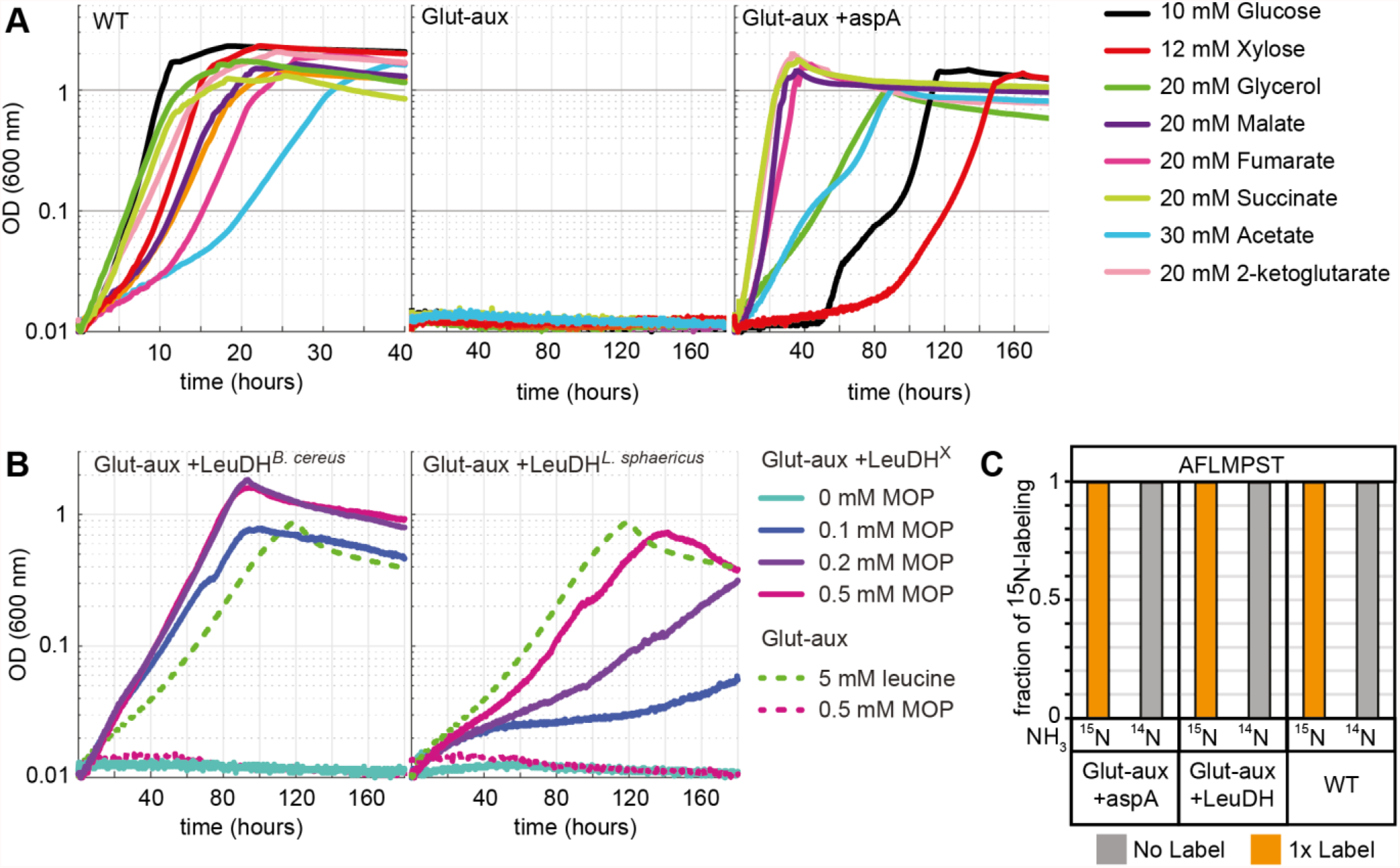
Aspartate ammonia lyase or leucine dehydrogenase can replace glutamate-based ammonia assimilation to fix all ammonium for cell growth. **A** Growth of WT, glut-aux and glut-aux +aspA on M9 with indicated carbon sources. Overexpression of aspartate ammonia lyase allows the glut-aux to assimilate free ammonium via fumarate amination. **B** Growth of glut-aux and glut-aux +LeuDH from *Bacillus cereus* or *Lysinibacillus sphaericus* on M9 with 20 mM glycerol as carbon source and the indicated additional substrates. Overexpression of LeuDH allows the glut-aux to assimilate free ammonium from the medium in presence of MOP. **C** Schematic representation of nitrogen labelling results for glut-aux +AspA, glut-aux +LeuDH and WT strains. Cells were grown either with ^15^NH_4_ or ^14^NH_4_ and 20mM succinate (glut-aux +AspA), 20 mM glycerol + 0.5 mM MOP (glutaux +LeuDH, from *B. cereus* or *L. sphaericus*) or 20 mM glycerol (WT). All strains were grown in M9 medium with ammonium. Data represents triplicate experiments with < 5 % variation.

### Leucine dehydrogenase

In our initial experiments, we showed that leucine can serve as amine group donor in the glut-aux strain (Fig. 2). To test if we can engineer the glut-aux strain to produce leucine for the concomitant transaminase reactions, we overexpressed leucine dehydrogenase, which catalyzes the reductive amination of 4-methyl-2-oxopentanoate (MOP) to leucine. Leucine dehydrogenase is absent in the repertoire of *E. coli’s* enzymes, hence we separately overexpressed two leucine dehydrogenase (LeuDH) genes, one from *Bacillus cereus* (Li et al, 2009) and the other from *Lysinibacillus sphaericus* (Lu et al, 2016). In initial experiments the glut-aux +LeuDH did not grow in minimal media with 20 mM glycerol. As MOP is not a central metabolite, its concentration might be limiting the activity of the enzyme. Consequently, we observed growth of the glut-aux strain only upon supplementation with catalytic amounts of MOP, which served as ammonium acceptor in the amination reaction catalyzed by leucine dehydrogenase (Fig 7B). We note that the MOP concentrations supplemented to the medium were at least 10-fold lower compared to the 5 mM leucine addition which served as a positive control in the experiments. The glut-aux strain overexpressing *Bacillus cereus* LeuDH growing on glycerol and 0.2- and 0.5-mM MOP reached twice the OD_600_ obtained for the glut-aux strain growing on glycerol and 5 mM leucine. This yield difference indicates a recycling of MOP through alternating activities of leucine dehydrogenase and transaminase, fixing ammonium and transferring amine groups to other ketoacids. Growth of the glut-aux strain overexpressing *L. sphaericus* LeuDH resulted in lower yields than achieved with *B. cereus* LeuDH overexpression, and also lower yields than seen for the glut-aux with 5 mM leucine, indicating lower enzyme efficiency under the tested conditions. To verify ammonium assimilation via leucine dehydrogenases the ^15^N-labeling in proteinogenic amino acids was analyzed after growing the glut-aux +LeuDH strains with 20 mM glycerol, 0.5 mM MOP, and either with ^15^NH_4_ or ^14^NH_4_ (Fig. 7C). The observed single label for the analyzed amino acids alanine, phenylalanine, methionine, proline, serine and threonine once again confirm operation of the leucine dehydrogenases ammonium entry point (Fig. 7C).

## Discussion

Our study provides a broad investigation on amino acid and ammonium metabolism in *E. coli*. To examine which amino acids can be used as amine source to compensate for the absence of canonical glutamate production, we created a glut-aux strain and tested its ability to grow on different amino acids. Surprisingly, the glut-aux did not grow on all tested amino acids. Also, some of the canonical transamination products of glutamate did not rescue growth of the glut-aux strain via the reverse reaction (i.e. alanine). For this amino acid, we identified alanine degradation as the counteracting pathway. The glut-aux strain was capable of growth on most amino acids (proline, aspartate, asparagine, methionine, leucine, isoleucine, valine, glutamate, glutamine, ornithine, tyrosine, phenylalanine, tryptophan and histidine). As absence of nitrogen from the medium only affected growth of the glut-aux on ornithine, we concluded that nitrogen response dependent regulation was of minor relevance for amino acid metabolism. We assumed that in order for the glut-aux strain to grow, these amino acids could either (i) fully replace glutamate as amine donor, (ii) could donate their amine directly to 2-ketoglutarate, (iii) were converted into an intermediate donating an amine to 2-ketoglutarate to form glutamate or (iv) were direct glutamate precursors. To investigate which reaction produced glutamate, we deleted several genes responsible for the respective amino acid degradation pathway. While proline was directly converted to glutamate (iv), utilization of histidine, tyrosine, phenylalanine, tryptophan, methionine, isoleucine and leucine depended on transamination mediated by AspC, one of the three main transaminases in *E. coli* (Gelfand & Steinberg, 1977). Three aspects of this finding were surprising to us. First, deletion of AspC alone was sufficient to abolish growth of the glut-aux strain on tyrosine, phenylalanine, isoleucine and leucine. This was unexpected, since the other two main transaminases TyrB and IlvE share cross-reactivities with these amino acids, and indicates that despite of their cross-reactivity they cannot fully replace AspC *in vivo*. Residual tyrosine and phenylalanine aminotransferase activity in tyrB and ilvE knockout strains in previous works led to the hypothesis that AspC was mainly exhibiting these activities, which herewith is further supported (Gelfand & Steinberg, 1977). Second, deletion of AspC abolished growth on histidine, methionine, isoleucine and leucine for which only low or no specific activity of AspC was reported *in vitro* (Hayashi et al, 1993; Powell & Morrison, 1978). Under physiological conditions, the substrate range of AspC thus seems to be broader than previously anticipated. Third, the glut-aux Δ*aspC* strain was unable to grow with tryptophan as amine source. Although activity of AspC with tryptophan was demonstrated *in vitro* (Hayashi et al, 1993; Powell & Morrison, 1978), transamination is not a known tryptophan degradation pathway in *E. coli* (Reitzer, 2014). We hence conclude that in the glut-aux strain, tryptophan is the direct substrate of AspC to form (indole-3-yl)-pyruvate and glutamate as in some bacteria (e.g. *Pseudomonas aeruginosa* (Bortolotti et al, 2016), *Clostridium sporogenes* (O’Neil & DeMoss, 1968)), protozoa (*Trichomonas vaginalis*, (Lowe & Rowe, 1985) or mammals (Shrawder & Martinez-Carrion, 1972). In summary, we expanded previous knowledge by several novel transamination routes in *E. coli*.

In addition to identifying natively present routes of amino acid metabolism, we demonstrated how amino acid metabolism can be rewired to allow utilization of new amine sources. Overexpression of alanine-2-oxoglutarate aminotransferase together with the removal of an alanine sink allowed usage of alanine as amine source. By overexpressing a transaminase transferring amines from β-alanine to 2-oxoglutarate we achieved usage of the non-proteinogenic amino acid β-alanine. Following the same logic, we deleted the D-amino acid dehydrogenase gene *dadA* as possible internal sink for glycine/serine and expressed an aspartate-glyoxylate aminotransferase to select for its reverse activity. The engineered strain was able to use the amine from glycine to support growth. Surprisingly, we discovered that the strain did not use glycine directly, but converted it to serine, which was yet another substrate of the transaminase, showing high activity for serine (*k*_cat_ of 25.87 s^-1^) with oxaloacetate as amine acceptor. In principle, the newly discovered serine-oxaloacetate transaminase activity can be coupled to formate assimilation via the reductive glycine pathway (Kim et al, 2020), which fixes ammonium by reverse activity of the glycine cleavage system. This allows ammonium transfer to make aspartate and convert serine to hydroxypyruvate instead of pyruvate (generated via serine deaminase activity in the reductive glycine pathway), hence saving some ATP which is needed to convert pyruvate into PEP, essential for anaplerosis and gluconeogenesis. As shown by the aforementioned examples, the glut-aux strain is an excellent selection platform to screen for any reactions producing amino acids which can rescue growth and by that allow use of new amine sources.

The function of glutamate as universal amine transfer molecule originates in its role as canonical entry point of ammonium into metabolism (Kumada et al, 1993). We were interested in finding out whether the prominence of this ammonium assimilation mechanism can be explained by a significant advantage compared to other mechanisms of ammonium assimilation. To investigate this, we replaced canonical ammonium assimilation via glutamate biosynthesis with two alternative pathways. Both, fumarate amination to aspartate and 4-methyl-2-oxopentanoate amination to leucine, reconstituted the ability of the glut-aux strain to assimilate free ammonium. Growth dependent on ammonium assimilation via AspA was fastest when a carbon source metabolically close to fumarate was provided. We note that assimilation of ammonium via aspartate ammonia-lyase is energetically different from ammonium assimilation via glutamate. While a glutamate dependent amination network requires NADPH consumption by glutamate dehydrogenase, an aspartate ammonia-lyase dependent network requires NADH for the reduction of oxaloacetate back to fumarate; oxaloacetate being the transamination product of aspC. NADH is energetically cheaper compared to NADPH, as the recovery of the latter from NADH wastes some proton motive force through the membrane bound transhydrogenase, and hence comes with the indirect cost of ATP synthesis (Spaans et al, 2015). Although ammonium assimilation via aspartate or leucine biosynthesis led to immediate growth of the glut-aux strain in growth-optimized laboratory conditions, these do not reflect natural conditions under which glutamate-based ammonium assimilation might have evolved and become prominent. Under ammonium limiting conditions, GDH and GS are kinetically superior with a *K*_M_ of 2 mM and 0.1 mM for ammonia, respectively (Reitzer, 2014) when compared to AspA or LeuDH from *Bacillus* cereus with *K*_M_ values of 20 mM and 13 mM for ammonia, respectively (Suzuki et al, 1973) (Sanwal & Zink, 1961). Additionally, the availability of two different systems (GDH and GS/GOGAT) which are each optimal for different growth conditions but generate the same molecule is unique about glutamate-based ammonium assimilation. The interplay of these systems allows a more flexible metabolic response to varying nitrogen and energy availability than leucine dehydrogenase or aspartate ammonia lyase and might thus explain the conservation of glutamate coupled ammonium assimilation, which is mirrored by presence of GS in all extant organisms (Kumada et al, 1993).

The possibility to modify ammonium metabolism by metabolic engineering indicates that the underlying metabolic network is highly flexible. Additionally, these findings might have relevance for the metabolic engineering of synthetic pathways, e.g. for growth coupled selection (Orsi et al, 2021) or the production of certain amino acids or their derivatives. By engineering ammonium assimilation via non-glutamate producing reactions, the cell’s need for ammonium assimilation can be decoupled from glutamate biosynthesis for the production of nitrogenous compounds. This view is in agreement with the paradigm of modular design for bioengineering, which can support new endeavors for strain design for biotechnological production processes. Here, the most obvious impact could be made on the million-ton scale industrial amino acid production (Wendisch, 2020). Therefore, engineering new ammonium entry points as well as extending the amination network could positively affect these processes.

Altogether, our research addressed three distinct and complementary aspects regarding amino acid and ammonium metabolism. We provided a comprehensive overview on the options and limitations of the cell’s amination network. Then, we showed its flexibility when engineering the network for new amine sources. Finally, by engineering new ammonium entry-points, we increased the potential design space for engineering ammonium assimilation and dissimilation.

## Methods

### Strains

All *E. coli* strains used in this study are listed in Table 1. Strain SIJ488, which carries inducible recombinase and flippase genes (Jensen et al, 2015), was used as wildtype for generation of deletions. Gene deletions were performed by λ-Red recombineering or P1-transduction as described below.

**Table 1.** Strains and plasmids used in this study.

| name | Description / deletions | References |
| --- | --- | --- |
| ***E. coli* strains** |  |  |
| DH5α | Cloning of overexpression constructs | |
| SIJ488 | WT, integrated λ-red recombinase and flippase | (Jensen et al, 2015) |
| glut-aux | $\Delta gdhA \Delta gltBD$ | This study |
| glut-aux $\Delta ybdL$ | $\Delta gdhA \Delta gltBD \Delta ybdL$ | This study |
| glut-aux $\Delta putA$ | $\Delta gdhA \Delta gltBD \Delta putA$ | This study |
| $\Delta putA$ | $\Delta putA$ | This study |
| glut-aux $\Delta aspC$ | $\Delta gdhA \Delta gltBD \Delta aspC$ | This study |
| glut-aux $\Delta dadX$ | $\Delta gdhA \Delta gltBD \Delta dadX$ | This study |
| glut-aux $\Delta dadA$ | $\Delta gdhA \Delta gltBD \Delta dadA$ | This study |
| glut-aux $\Delta gcv$ | $\Delta gdhA \Delta gltBD \Delta gcvTHP$ | This study |
| glut-aux $\Delta gcv \Delta dadA$ | $\Delta gdhA \Delta gltBD \Delta gcvTHP \Delta dadA$ | This study |
| JW0999 | KEIO $\Delta putA$ | (Baba et al, 2006) |
| **Plasmids** |  |  |
| ASS | Over-expression plasmid with p15A origin, Streptomycin resistance, constitutive strong promoter | (Wenk et al, 2018) |
| ASS-aspA | ASS backbone for overexpression of *aspA* from *E. coli* | This study |
| ASS-alaA | ASS backbone for overexpression of *alaA* from *E. coli* | This study |
| ASS-alaC | ASS backbone for overexpression of *alaC* from *E. coli* | This study |
| ASS-leuDH<sup>B. cereus</sup> | ASS backbone for overexpression of leucine dehydrogenase from *Bacillus cereus* | This study |
| ASS-leuDH<sup>L. sphaericus</sup> | ASS backbone for overexpression of leucine dehydrogenase from *Lysinibacillus sphaericus* | This study |
| ASS-BAPAT | ASS backbone for overexpression of β-alanine-pyruvate-aminotransferase from *Pseudomonas aeruginosa* | This study |
| ASS-BAOAT | ASS backbone for overexpression of β-alanine-2-oxoglutarate-aminotransferase from *Saccharomyces kluyveri* | This study |
| ASS-BhcA | ASS backbone for overexpression of glycine-oxaloacetate aminotransferase from *Paracoccus denitrificans* | This study |

### Gene deletion via P1 transduction

Deletions of *putA* and *dadX* were generated by P1 phage transduction (Thomason et al, 2007). Strains from the Keio collection carrying single gene deletions with a kanamycin-resistance gene (KmR) as selective marker were used as donor strains (Baba et al, 2006). Selection for strains that had acquired the desired deletion was performed by plating on appropriate antibiotics (Kanamycin, Km) and confirmed by determining the size of the respective genomic locus *via* PCR using DreamTaq polymerase (Thermo Scientific, Dreieich, Germany) and the respective KO-Ver primers (Supplementary Table S4). Additionally, it was confirmed that no copy of the gene to be deleted was present anywhere in the genome by PCR using DreamTaq polymerase (Thermo Scientific, Dreieich, Germany) and internal primers binding inside of the coding sequence of the gene. To remove the selective marker, a fresh culture was grown to OD_600_ ∼ 0.2, followed by inducing flippase expression by adding 50mM L-rhamnose and cultivating for ∼4h at 30°C. Loss of the antibiotic resistance was confirmed by identifying individual colonies that only grew on LB in absence of the respective antibiotic and by PCR of the genomic locus using the locus specific KO-Ver primers.

### Gene deletion by recombineering

For gene deletion by recombineering, kanamycin resistance cassettes were generated via PCR – ‘KO’ primers with 50 bp homologous arms are listed in Supplementary Table S4 – using the kanamycin (Km) cassette from pKD4 (pKD4 was a gift from Barry L. Wanner (Addgene plasmid # 45605; http://n2t.net/addgene:45605; RRID:Addgene_45605), (Datsenko & Wanner, 2000)) ((Baba et al, 2006)), or in case of the *aspC* deletion the chloramphenicol (Cap) cassette from pKD3 (pKD3 was a gift from Barry L. Wanner (Addgene plasmid # 45604; http://n2t.net/addgene:45604; RRID:Addgene_45604)). To prepare cells for gene deletion, fresh cultures were inoculated in LB and the recombinase genes were induced by addition of 15 mM L-arabinose at OD ∼0.4-0.5. After incubation for 45 min at 37°C cells were harvested and washed three times with ice cold 10 % glycerol (11,000 rpm, 30 sec, 2°C). ∼300 ng of Km cassette PCR-product was transformed via electroporation (1 mm cuvette, 1.8 kV, 25 μF, 200 Ω). After selection on kanamycin, gene deletions were confirmed via PCR using ‘KO-Ver’ primers (Supplementary Table S4). To remove the Km cassette, 50 mM L-rhamnose, which induces flippase gene expression, was added to an exponentially growing 2 ml LB culture at OD 0.5; induction time was ≥ 3 h at 30°C. Colonies were screened for kanamycin sensitivity and removal of antibiotic resistance cassette was confirmed by PCR (using ‘KO-Ver’ primers).

### Plasmid construction

For overexpression, genes encoding for leucine dehydrogenases from *Bacillus cereus* (LeuDH^*B*. *cereus*^, P0A392) and *Lysinibacillus sphaericus* (LeuDH^*L*. *sphaericus*^, B1HRW1), aspartate ammonia-lyase from *E. coli* (AspA, P0AC38), β-alanine oxoglutarate transaminase from *Saccharomyces kluyveri* (BAOAT, A5H0J5), β-alanine pyruvate transaminase from *Pseudomonas aeruginosa* (BAPAT, Q9I700), and Glycine-oxaloacetate-aminotransferase from *Paracoccus denitrificans* (BhcA, A1B8Z3) were synthesized after removal of restriction sites relevant for cloning (Zelcbuch et al, 2013) and codon adaptation to *E. coli’*s codon usage (Grote et al, 2005). Genes were synthesized by Twist Bioscience (San Francisco, CA, USA). Alanine transaminase genes *alaA* and *alaC* were amplified from *E. coli*’s genome with high-fidelity Phusion Polymerase (Thermo Scientific, Dreieich, Germany) using primer pairs *alaA*-amp_fwd, *alaA*-amp_rvs and *alaC*-amp_fwd, *alaC*-amp_rvs, respectively (Supplementary Table S4).

Cloning was carried out in *E. coli* DH5α. All genes were cloned via *Mph1103*I and *Xho*I into pNivC vector downstream of ribosome binding site “C” (AAGTTAAGAGGCAAGA) (Zelcbuch et al, 2013). Restriction enzymes *EcoR*I and *Pst*I (FastDigest, Thermo Scientific) were used to transfer the genes into the expression vector pZ-ASS (p15A origin, Streptomycin resistance, strong promoter) (Braatsch et al, 2008). Constructed vectors were confirmed by Sanger sequencing (LGC Genomics, Berlin, DE). The software Geneious 8 (Biomatters, New Zealand) was used for *in silico* cloning and sequence analysis.

The plasmid for the expression of the *dadA* gene was retrieved from the ASKA collection (Kitagawa et al, 2005). The plasmids for the expression of the *bhcA* and *ghrA* genes were described in previous publications (Miller et al, 2020; Schada von Borzyskowski et al, 2019).

### Media and growth experiments

LB medium (1% NaCl, 0.5% yeast extract, 1% tryptone) was used for cloning, generation of deletion strains, and strain maintenance. When appropriate, kanamycin (25 μg/mL), ampicillin (100 μg/mL), streptomycin, (100 μg/mL), or chloramphenicol (30 μg/mL) were used but omitted for growth experiments. Growth experiments were carried out in standard M9 minimal media (50 mM Na_2_HPO_4_, 20 mM KH_2_PO_4_, 1 mM NaCl, 20 mM NH_4_Cl, 2 mM MgSO_4_ and 100 μM CaCl_2_, 134 μM EDTA, 13 μM FeCl_3_·6H_2_O, 6.2 μM ZnCl_2_, 0.76 μM CuCl_2_·2H_2_O, 0.42 μM CoCl_2_·2H_2_O, 1.62 μM H_3_BO_3_, 0.081 μM MnCl_2_·4H_2_O) or in M9 media lacking NH_4_Cl. Carbon sources were used as indicated in the text. Unless otherwise specified, L-isomers of amino acids were used if extant. For growth experiments overnight cultures were incubated in 4 mL M9 medium containing 20 mM glycerol supplemented with 5 mM aspartate. Cultures were harvested (6,000\**g*, 3 min) and washed three times in M9 medium (w/ or w/o NH_4_Cl) to remove residual carbon and NH_4_Cl sources. Washed cells were used to inoculate growth experiments to an optical density (OD_600_) of 0.01 in 96-well microtiter plates (Nunclon Delta Surface, Thermo Scientific) at 37°C. Each well contained 150 μL of culture and 50 μL mineral oil (Sigma-Aldrich) to avoid evaporation while allowing gas exchange. Growth was monitored in technical triplicates at 37°C in BioTek Epoch 2 Microtiterplate reader (BioTek, Bad Friedrichshall, Germany) by absorbance measurements (OD_600_) of each well every ∼10 minutes with intermittent orbital and linear shaking. Blank measurements were subtracted and OD_600_ measurements were converted to cuvette OD_600_ values by multiplying with a factor of 4.35, as previously established empirically for the instruments. Growth curves were plotted in MATLAB and represent averages of measurements of technical replicates.

### Isolation and sequence analysis of glut-aux ASS-*alaA* mutants

The glut-aux strain +*alaA* was inoculated to OD_600_ of 0.02 in tube cultures of 4 mL M9 + 20mM glycerol + 5mM alanine. Cell growth was monitored during prolonged incubation at 37°C for 7-14 days. Within that time several cultures started to grow and reached an OD above 1.0. Cells were streaked out on LB plates with streptomycin (to maintain the pZ-ASS-*alaA* plasmid) by dilution streak to generate single colonies. Isolates were inoculated into tube cultures of 4 mL M9 + 20mM glycerol + 5mM alanine, and the ones which immediately grew were used in genome sequence analysis. Genomic DNA was extracted using the Macherey-Nagel NucleoSpin Microbial DNA purification Kit (Macherey-Nagel, Düren, Germany) from 2×10^9^ cells of an overnight culture in LB medium supplied with streptomycin and chloramphenicol (to maintain pZ-ASS-*alaA* plasmid). Construction of (microbial short insert libraries) PCR-free libraries for single-nucleotide variant detection and generation of 150 bp paired-end reads on an Illumina HiSeq 3000 platform were performed by Novogene (Cambridge, UK). Reads were mapped to the reference genome of *E*.*coli* MG1655 (GenBank accession no. U00096.3) using the software Breseq (Barrick Lab, Texas) (Deatherage & Barrick, 2014). Using algorithms supplied by the software package, we identified single-nucleotide variants (with >50% prevalence in all mapped reads) and searched for regions with coverage deviating more than 2 standard deviations from the global median coverage.

### ^15^N isotopic labelling of proteinogenic amino acids

To elucidate the origin of the nitrogen in amino acids we used ^15^N isotope tracing experiments. Proteinogenic amino acids were analyzed after cell growth in M9 containing ^15^NH_4_Cl (Sigma-Aldrich, Germany). Cells (1 mL of OD_600_ 1) were harvested by centrifugation (6,000*g) after reaching stationary phase and washed in H_2_O. Proteins were hydrolyzed in 6 N HCl, at 95°C for 24 h (You et al, 2012). HCl was removed by evaporation under an air stream at 95°C. Samples were then resuspended in 1 ml H2O, insoluble compounds were removed by centrifugation (10 min, 16,000*g), and supernatants were used for analysis. Amino acid masses were analyzed by UPLC-ESI-MS as described previously (Giavalisco et al, 2011) with a Waters Acquity UPLC system (Waters) using a HSS T3 C18 reversed phase column (100 mm × 2.1 mm, 1.8 μm; Waters). The mobile phases were 0.1 % formic acid in H2O (A) and 0.1% formic acid in acetonitrile (B). The flow rate was 0.4 mL/min with a gradient of 0 to 1 min – 99% A; 1 to 5 min – linear gradient from 99% A to 82%; 5 to 6 min – linear gradient from 82% A to 1% A; 6 to 8 min – kept at 1% A; 8-8.5 min – linear gradient to 99% A; 8.5-11 min – re-equilibrate. Mass spectra were acquired using an Exactive mass spectrometer (Thermo Scientific) in positive ionization mode, with a scan range of 50.0 to 300.0 m/z. The spectra were recorded during the first 5 min of the LC gradients. Data analysis was performed using Xcalibur (Thermo Scientific). Amino acid standards (Sigma-Aldrich, Germany) were analyzed for determination of the retention times under the same conditions.

### Protein purification

To produce DadA and GhrA proteins for *in vitro* characterization, *E. coli* BL21 cells were transformed with the plasmid containing the respective gene. The cells were then grown on LB agar plates containing 50 μg/mL kanamycin or 100 μg/mL ampicillin at 37 °C overnight. Grown cells were used to inoculate a liter of selective terrific broth (TB). The expression cultures were grown overnight at 25 °C in a shaking incubator. Their biomass was collected by centrifugation at 6,000 g for 15 min at room temperature. The cells were resuspended in twice their volume of buffer A (50 mM HEPES/KOH pH 7.8, 500 mM NaCl). The cells were lysed with a Microfluidizer (LM-10 H10Z, Microfludics, Westwood, US) at 16.000 PSI for three passes on ice and the lysate was cleared by ultracentrifugation at 100,000×g for 45 min at 4 °C and subsequently filtered through a 0.45 μm PTFE filter. The filtered lysate was loaded onto a 1 ml HisTrap FF (GE Healthcare, Freiburg, Germany) and unbound protein was removed with 20 column volumes of 15% buffer B (50 mM HEPES/KOH pH 7.8, 500 mM NaCl, 500 mM Imidazole) in buffer A. The protein was then eluted in 100% buffer B. The protein was desalted with a HiTrap 5 ml Desalting column (GE Healthcare, Freiburg, Germany) and a desalting buffer (50 mM HEPES/KOH, 50 mM NaCl, 20% (v/v) Glycerol). Protein concentrations were determined by the protein’s theoretical extinction coefficient and their absorbance at 280 nm. BhcA was expressed and purified as described previously (Schada von Borzyskowski et al, 2019).

### Measurement of enzyme activity

The activities of all tested enzymes were measured with a Cary 60 UV-Vis spectrophotometer (Agilent Technologies GmbH, Waldbronn, Germany) at 37°C. To test the transaminase activity of BhcA with serine, the absorbance at 365 nm of the reaction mix (50 mM HEPES/KOH pH 7.5, 0.7 mM NADPH, 7 mM OAA, 30 μg GhrA, and 2.5 μg BhcA; modified based on Schada von Borzyskowski et al., 2019) was tracked over time. The reaction was started by adding varying concentrations of serine to the mixture and the resulting slope in absorbance decrease was measured. To test the activity of BhcA for oxaloacetate, another reaction mix (50 mM HEPES/KOH pH 7.5, 0.7 mM NADPH, 100 mM serine, 30 μg GhrA, and 2.5 μg BhcA) was prepared. This time the reaction was started by adding oxaloacetate in varying concentrations to the mix. Activities of DadA were measured via following the absorbance of Dichlorophenolindophenole (DCPIP) at 600 nm. To measure activity with D-alanine the reaction mix (200 mM HEPES/KOH pH 7.5, 0.1mM DCPIP, 1.5 mM PES, 10 mM KCN, 60 μg DadA) was started with varying concentrations of D-alanine. To measure the same reaction for glycine, the amount of DadA was increased to 280 μg, and the reaction was started with glycine.

## Acknowledgements

This work was funded by the Max Planck Society. S.N.L. acknowledges support from the BMBF grant ForceYield (031B0825B). T.J.E. acknowledges additional support from the German Research Foundation (SFB987 ‘Microbial diversity in environmental signal response’).

## Author contributions

S.N.L. and A.B.-E. conceived and supervised the study. S.N.L., A.B.-E. and S.A. designed the experiments. H.S.M., A.M. and T.W. constructed plasmids and strains and performed growth experiments. H.S.M. analyzed the genome sequencing data. H.S.M. performed and analyzed ^15^N-labeling experiments, P.P. and L.S.v.B. purified enzymes and analyzed enzyme activities. S.N.L., H.S.M, and T.J.E. analyzed the results and wrote the manuscript with contributions from all authors.

## Supplementary material

**Figure S1:**
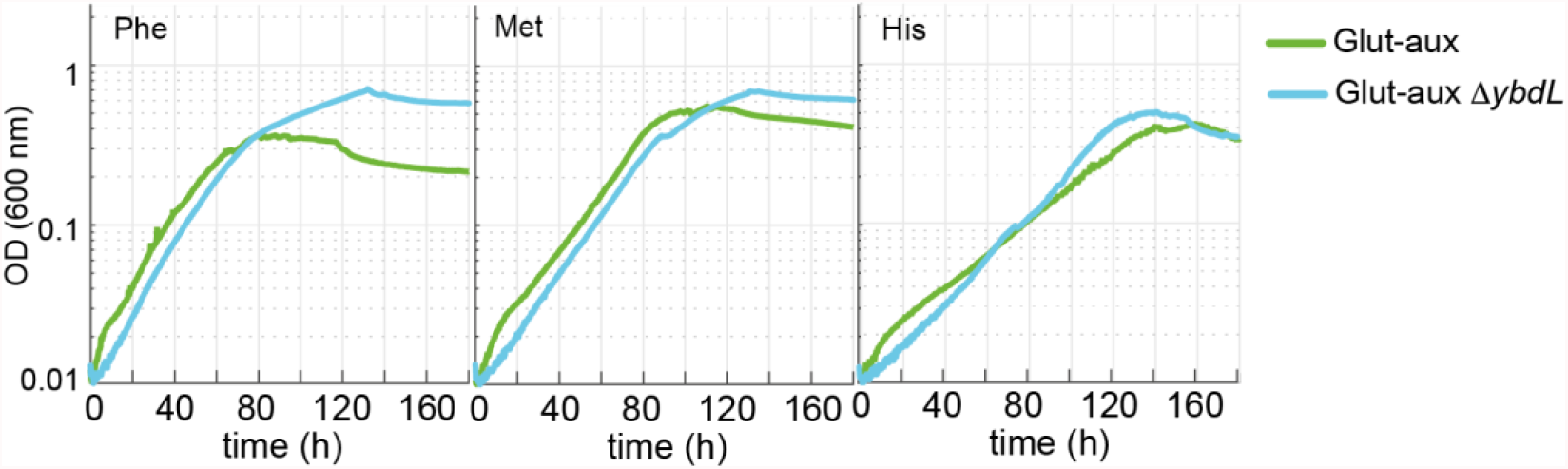
Deletion of *ybdL* does not alter growth of the glut-aux strain with phenylalanine, methionine and histidine as amine sources. Experiments were carried out in M9 w/ ammonium containing 20 mM glycerol and 5 mM of the indicated amine-sources. Data represents means of triplicates with < 5 % variation.

**Figure S2:**
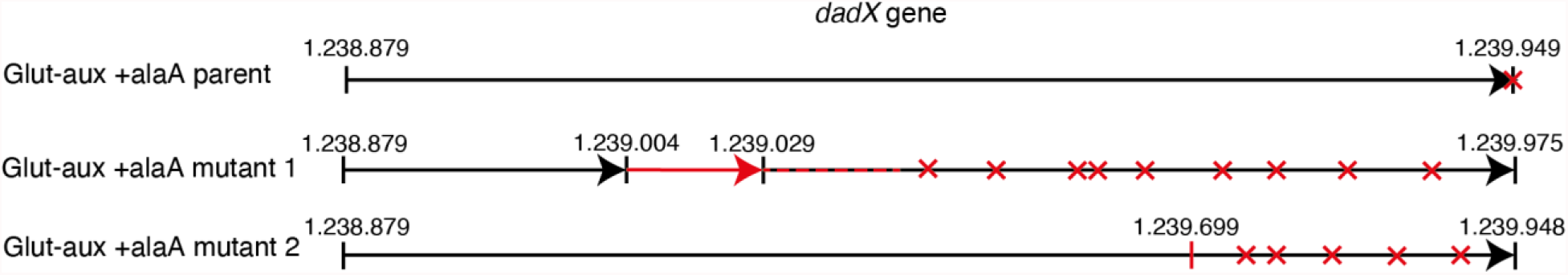
Schematic presentation of 26 bp duplication (red) found in the *dadX* gene of the glut-aux +alaA mutant 1 and 1 bp deletion in the dadX gene glut-aux +AlaA mutants 2 and 3 in comparison to the glut-aux +alaA parent. Both mutations cause stop codons (red crosses) within the *dadX* gene reading frame.

**Figure S3:**
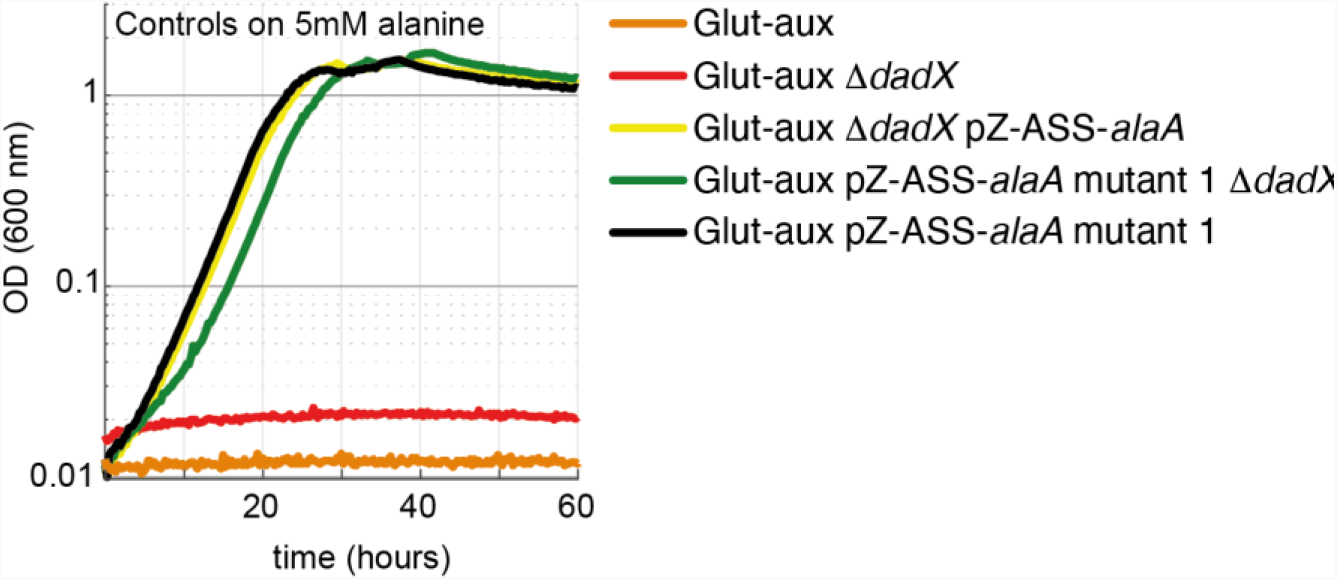
DadX mutation or deletion and alanine transaminase overexpression allows alanine utilization as amine source. Growth of the glut-aux strains on M9 w/ammonium and 20 mM glycerol with 5 mM alanine as amine donor. Strains not overexpressing alanine transaminase cannot utilize alanine as amine source. All data represents triplicate measurements with < 5 % variation.

**Figure S4:**
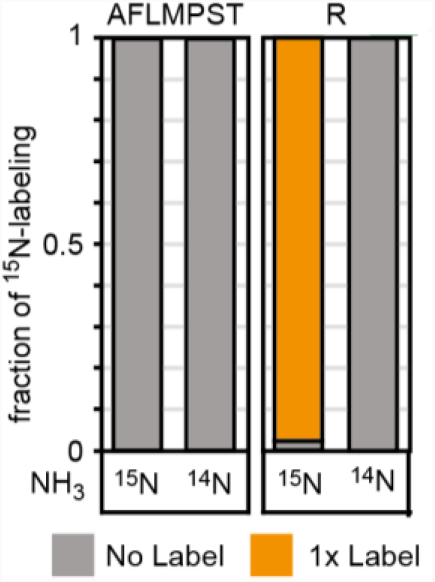
^15^N-labelling experiments confirm incorporation of the amino group of β-alanine into proteinogenic amino acids (single letter code). The glut-aux +BaOAT strain was grown in medium containing 20 mM glycerol with 5 mM β-alanine and either ^14^NH_4_ or ^15^NH_4_. Data represents means of three independent experiments.

**Figure S5:**
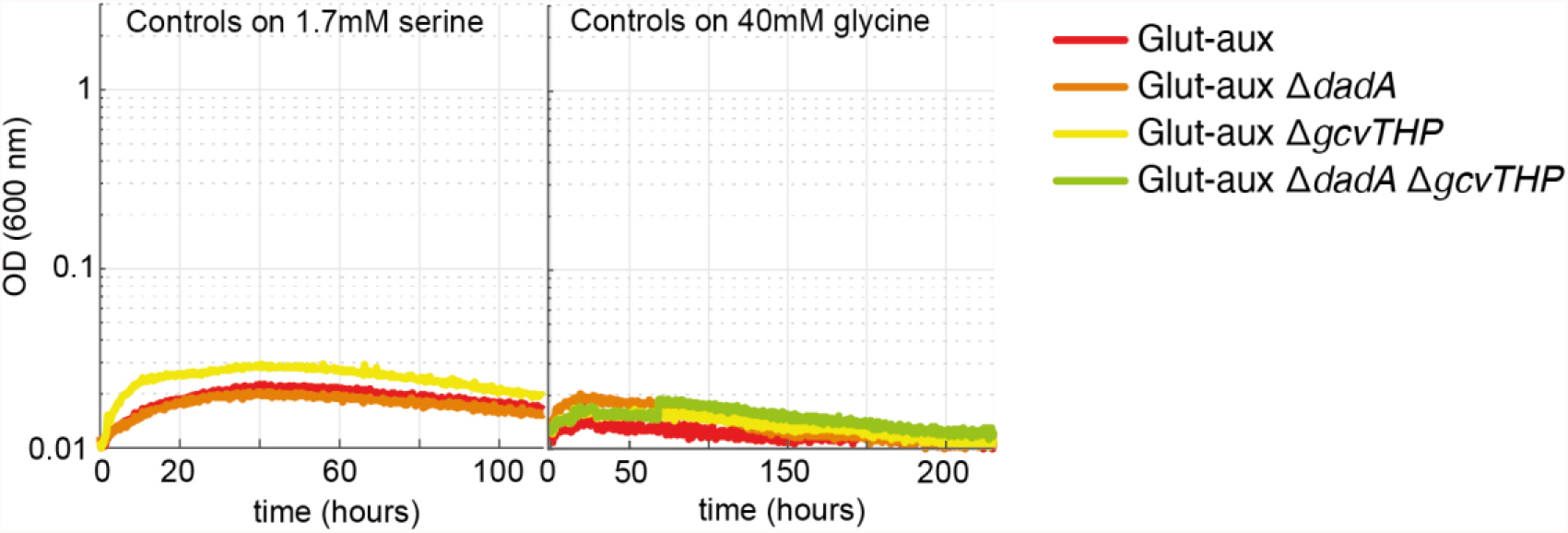
Transaminase overexpression is essential for use of serine or glycine as amine donor in the glut-aux strain. Glut-aux strains not overexpressing BhcA were incubated in M9 w/ ammonium and 20 mM glycerol and 1.7 mM serine or 40 mM glycine as amine donor. No growth was obtained, indicating the need for BhcA overexpression for utilization of glycine or serine as amine sources. All data represents triplicate measurements with < 5 % variation.

**Figure S6:**
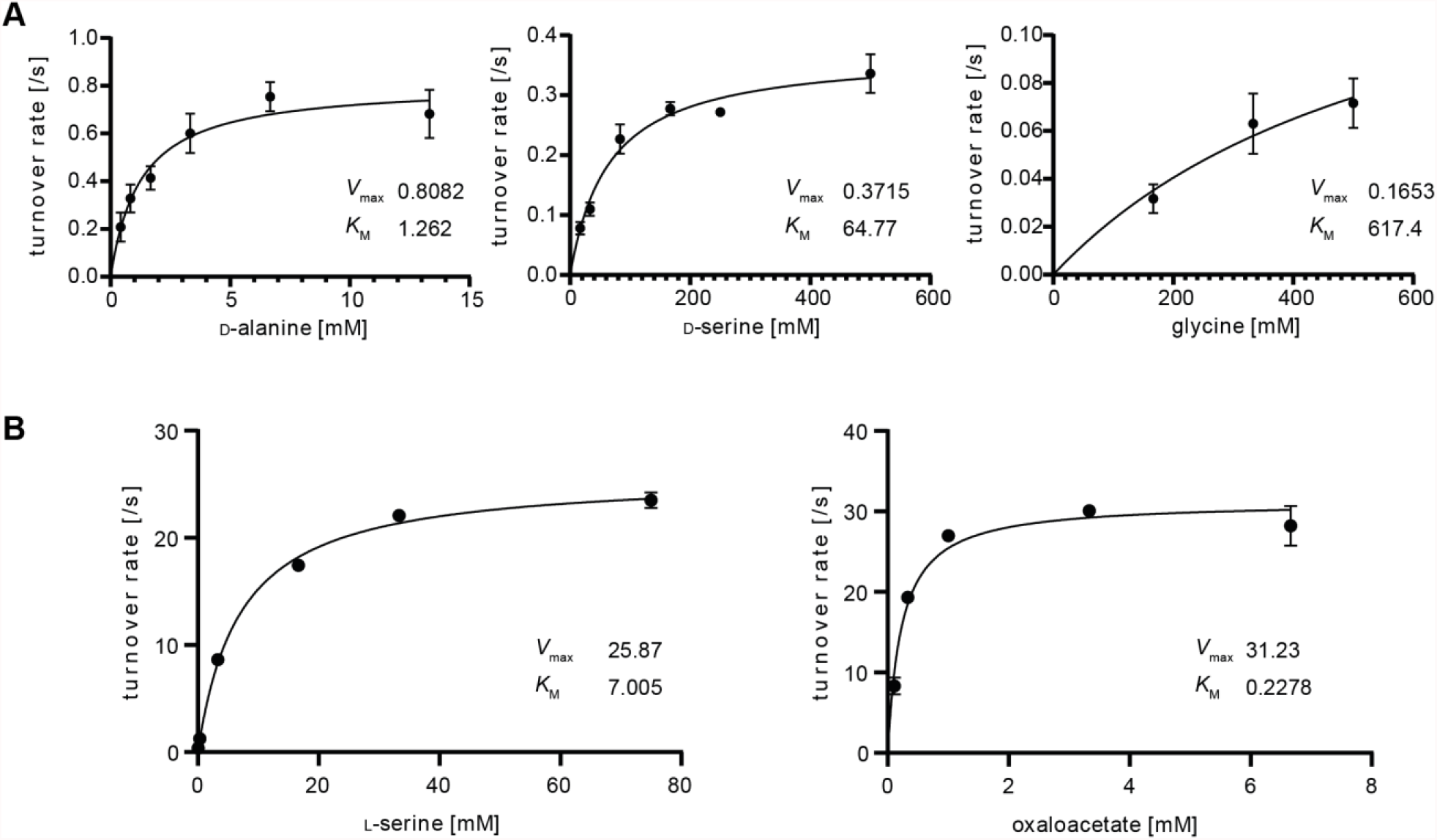
Michaelis-Menten kinetics of DadA and BhcA for selected substrates. **A** DadA turnover rate with D-alanine (left), D-serine (middle) and glycine (right) measured in a DCPIP-coupled assay. **B** BhcA turnover rate with serine (left) and oxaloacetate (right). To saturating concentrations of oxaloacetate (left) or serine (right), different concentrations of the respective other substrate were added in an assay coupling BhcA mediated hydroxypyruvate formation to NADPH dependent hydroxypyruvate reduction catalyzed by GhrA. Data are shown from n = 3 independent experiments at different substrate concentrations. Kinetic parameters are listed in Supplementary Table 2 (DadA) and Supplementary Table 3 (BhcA).

**Figure S7:**
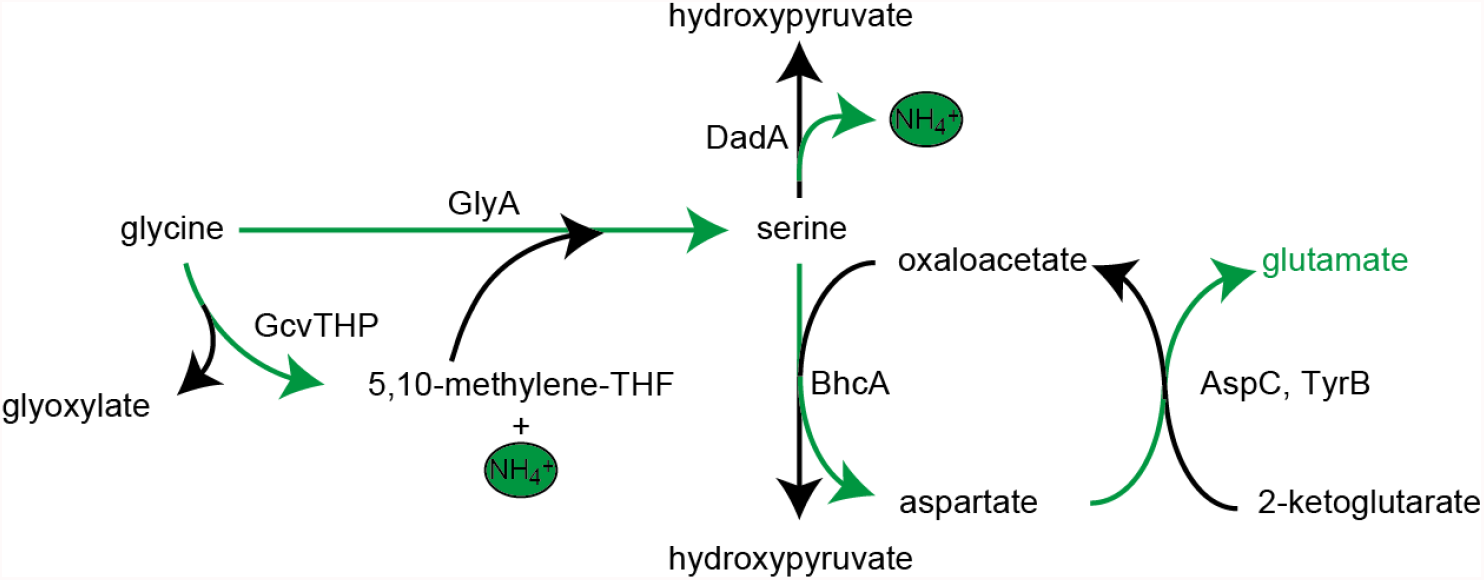
Schematic presentation of glycine and serine utilization by the glut-aux strain Δ*dadA* expressing BhcA. Glycine is converted to serine by combined activity of GlyA and GcvTHP, which is then substrate for BhcA in a transamination reaction with oxaloacetate. The formed aspartate allows glutamate formation via native transaminases. DadA promiscuously deaminating serine to form hydroxypyruvate might be an intracellular serine sink and needed to be deleted for optimal growth of the glutaux +BhcA with glycine or serine as amine source. Notably, the BhcA substrate L-serine would first need to be converted to D-serine, for example by promiscuously acting alanine racemase Alr, to be substrate for DadA. Green arrows indicate the transfer of ammonium.

**Figure S8:**
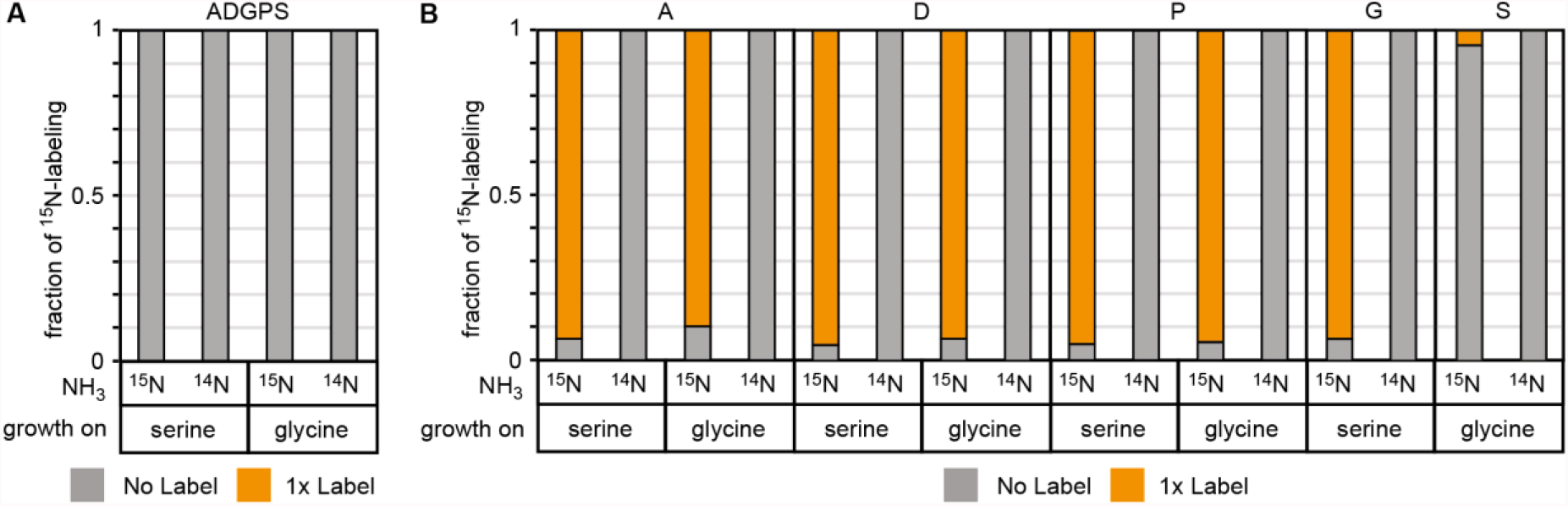
^15^N-labeling confirms amine donation from serine or glycine in glut-aux Δ*dadA* +BhcA. **A** Glut-aux Δ*dadA* +BhcA grown on ^15^N-NH_3_Cl M9 medium or ^14^N-NH_3_Cl M9 medium with 20 mM glycerol as carbon source. **B** WT grown on ^15^N-NH_3_Cl M9 medium or ^14^N-NH_3_Cl M9 medium with 20 mM glycerol as carbon source. For both strains, ^15^N labeling in amino acids (single letter code) was analyzed upon feeding with 1.7 mM of unlabeled serine or 20 mM of unlabeled glycine. Data represents means of triplicate measurements with errors < 5 %.

**Table S1.**
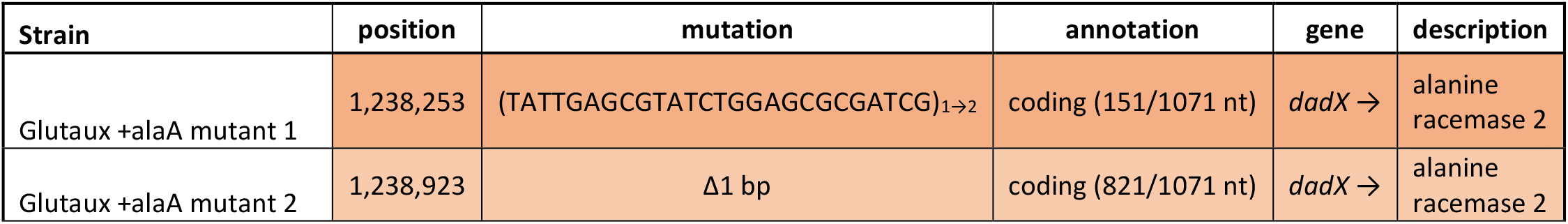
Mutations found in Glutaux +alaA mutants 1 and 2. After sequencing the genomes of two independently isolated glutaux +alaA mutants and the glutaux +alaA parent, the results were mapped against the *E. coli* MG1655 reference genome (GenBank accession no. U00096.3) using breseq. Mutations occurring in the mutants and not the parent are listed.

**Table S2.** Kinetics of DadA with D-alanine and glycine. Enzyme activity was determined in a DCPIP-coupled assay with various substrate concentrations and 60 μg protein for measurements with D-alanine and 280 μg protein for measurements with glycine. Data are mean ± SE.

| substrate | DadA |  |  |
| --- | --- | --- | --- |
| | $k_{cat}$ (s<sup>-1</sup>) | $K_M$ (mM) | $k_{cat}/K_M$ (M<sup>-1</sup> s<sup>-1</sup>) |
| D-alanine | 0.81 ± 0.05 | 1.26 ± 0.26 | 6.42 × 10<sup>2</sup> |
| D-serine | 0.37 ± 0.02 | 64.77 ± 9.28 | 5.71 × 10<sup>0</sup> |
| glycine | 0.17 ± 0.08 | 617.40 ± 463.2 | 2.75 × 10<sup>-1</sup> |

**Table S3.** Kinetics of BhcA with L-serine and oxaloacetate. Enzyme activity was determined in an assay coupling BhcA dependent L-serine transamination to NADPH dependent hydroxypyruvate reduction by GhrA with varying concentrations of L-serine or oxaloacetate for the respective activity measurements. Data are mean ± SE.

|  | BhcA |  |  |
| --- | --- | --- | --- |
| substrate | $k_{cat}$ ($s^{-1}$) | $K_M$ (mM) | $k_{cat}/K_M$ ($M^{-1} s^{-1}$) |
| L-serine | $25.87 \pm 0.4$ | $7.01 \pm 0.46$ | $3.7 \times 10^3$ |
| oxaloacetate | $31.23 \pm 0.89$ | $0.23 \pm 0.03$ | $1.4 \times 10^5$ |

**Table S4.**
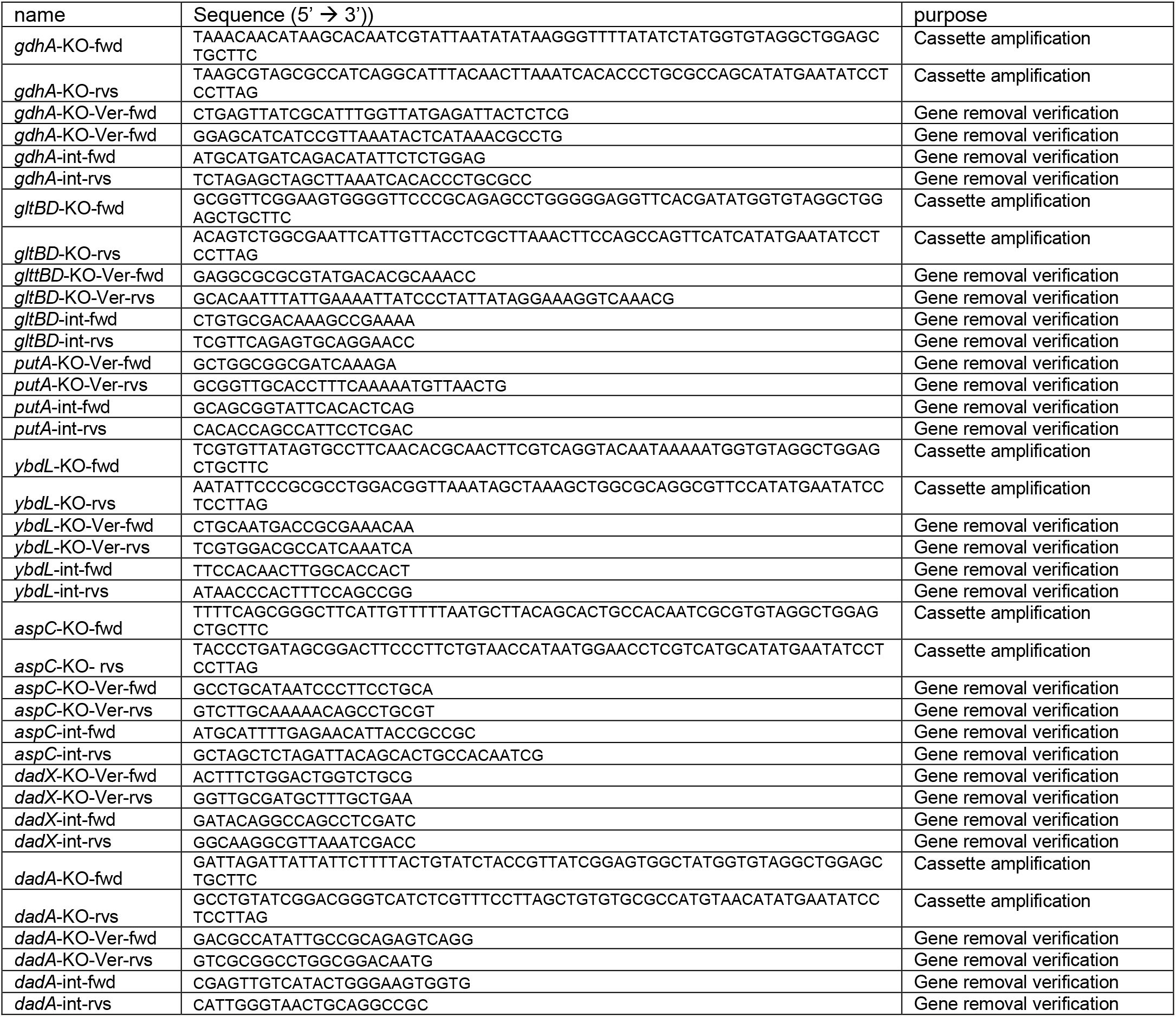
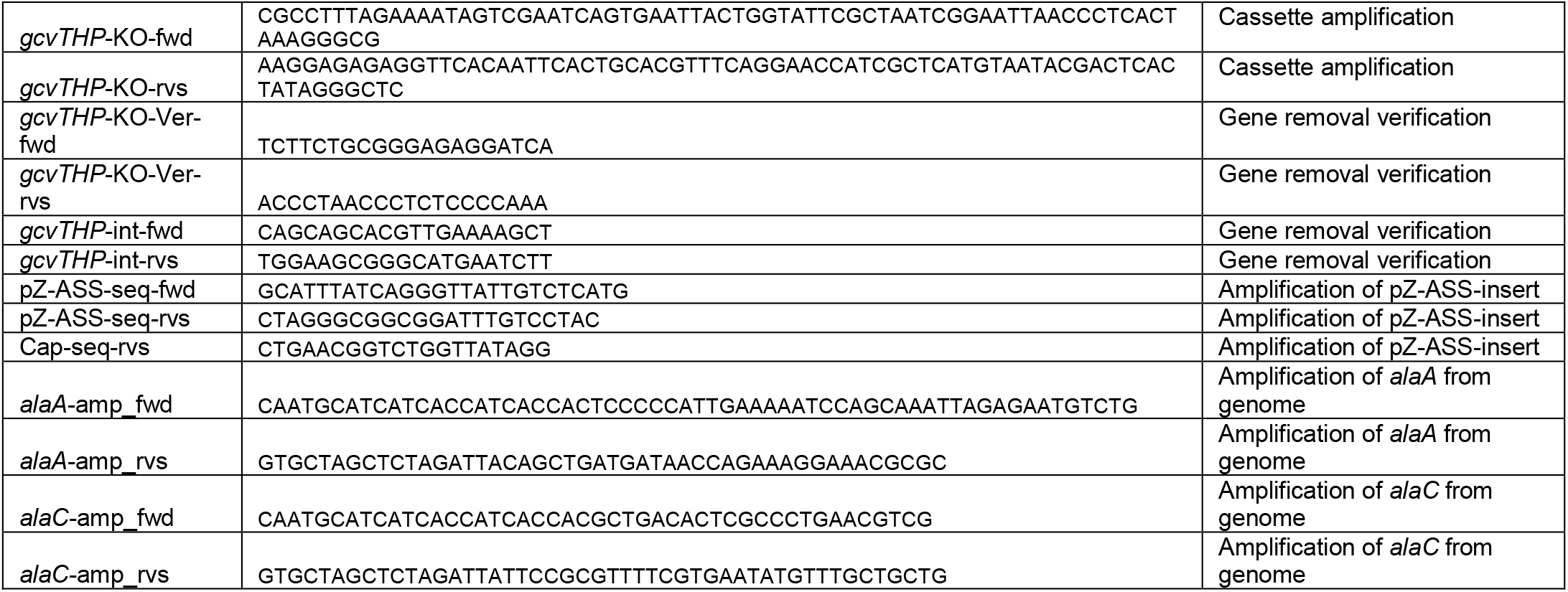
Oligonucleotide primers used. ‘KO’ primers were used to amplify the knockout Km cassette from pKD4 with 50 bp gene-specific upstream and downstream sequences. ‘KO-Ver’-primers (knockout-verification) were used to verify gene replacement by kanamycin resistance cassette and cassette removal by flippase. External and internal primers were used to verify successful removal of the gene from the genome.

## Appendix

### Sequences of synthetic genes

#### aspA

ATGCATCATCACCATCACCACTCTAACAACATCCGTATCGAAGAAGACCTGCTGGGTACCCGTGAAGTTCCGGCTGACGCTTACTACGGTGTTCACACCCTGCGTGCTATCGAAAACTTCTACATCTCTAACAACAAAATCTCTGACATCCCGGAATTTGTTCGTGGTATGGTTATGGTTAAAAAAGCTGCTGCTATGGCTAACAAAGAACTGCAAACCATCCCGAAATCTGTTGCTAACGCTATCATCGCTGCTTGCGACGAAGTTCTGAACAACGGTAAATGTATGGACCAGTTCCCGGTTGACGTTTACCAGGGTGGTGCTGGTACCTCTGTTAACATGAACACCAACGAAGTTCTGGCTAACATCGGTCTGGAACTGATGGGTCACCAGAAAGGTGAATACCAGTACCTGAACCCGAACGACCACGTTAACAAATGCCAGTCTACCAACGACGCTTACCCGACCGGTTTCCGTATCGCTGTTTACTCTTCTCTGATCAAACTGGTTGACGCTATCAACCAGCTGCGTGAAGGTTTCGAACGTAAAGCTGTTGAATTTCAGGACATCCTGAAAATGGGTCGTACCCAGCTGCAAGACGCTGTTCCGATGACCCTGGGTCAGGAATTTCGTGCTTTCTCTATCCTGCTGAAAGAAGAAGTTAAAAACATCCAGCGTACCGCTGAACTGCTGCTGGAAGTTAACCTGGGTGCTACCGCTATCGGTACCGGTCTGAACACCCCGAAAGAATACTCTCCGCTGGCTGTTAAAAAACTGGCTGAAGTTACCGGTTTCCCGTGCGTTCCGGCTGAAGACCTGATCGAAGCTACCTCTGACTGCGGTGCTTACGTTATGGTTCACGGTGCTCTGAAACGTCTGGCTGTTAAAATGTCTAAAATCTGCAACGACCTGCGTCTGCTGTCTTCTGGTCCGCGTGCTGGTCTGAACGAAATCAACCTGCCGGAACTGCAAGCTGGTTCTTCTATCATGCCGGCTAAAGTTAACCCGGTTGTTCCGGAAGTTGTTAACCAGGTTTGCTTCAAAGTTATCGGTAACGACACCACCGTTACCATGGCTGCTGAAGCTGGTCAGCTGCAACTGAACGTTATGGAACCGGTTATCGGTCAGGCTATGTTCGAATCTGTTCACATCCTGACCAACGCTTGCTACAACCTGCTGGAAAAATGTATCAACGGTATCACCGCTAACAAAGAAGTTTGCGAAGGTTACGTTTACAACTCTATCGGTATCGTTACCTACCTGAACCCGTTCATCGGTCACCACAACGGTGACATCGTTGGTAAAATCTGCGCTGAAACCGGTAAATCTGTTCGTGAAGTTGTTCTGGAACGTGGTCTGCTGACCGAAGCTGAACTGGACGACATCTTCTCTGTTCAGAACCTGATGCACCCGGCTTACAAAGCTAAACGTTACACCGACGAATCTGAACAGTAATCTAGAGCTAGCG

#### BaOAT

ATGCATCATCACCATCACCACCCGTCTTACTCTGTTGCTGAACTGTACTACCCGGACGAACCGACCGAACCGAAAATCTCTACCTCTTCTTACCCGGGTCCGAAAGCTAAACAGGAACTGGAAAAACTGTCTAACGTTTTCGACACCCGTGCTGCTTACCTGCTGGCTGACTACTACAAATCTCGTGGTAACTACATCGTTGACCAGGACGGTAACGTTCTGCTGGACGTTTACGCTCAGATCTCTTCTATCGCTCTGGGTTACAACAACCCGGAAATCCTGAAAGTTGCTAAATCTGACGCTATGTCTGTTGCTCTGGCTAACCGTCCGGCTCTGGCTTGCTTCCCGTCTAACGACTACGGTCAGCTGCTGGAAGACGGTCTGCTGAAAGCTGCTCCGCAGGGTCAGGACAAAATCTGGACCGCTCTGTCTGGTTCTGACGCTAACGAAACCGCTTTCAAAGCTTGCTTCATGTACCAGGCTGCTAAAAAACGTAACGGTCGTTCTTTCTCTACCGAAGAACTGGAATCTGTTATGGACAACCAGCTGCCGGGTACCTCTGAAATGGTTATCTGCTCTTTCGAAAAAGGTTTCCACGGTCGTCTGTTCGGTTCTCTGTCTACCACCCGTTCTAAACCGATCCACAAACTGGACATCCCGGCTTTCAACTGGCCGAAAGCTCC GTTCCCGGACCTGAAATACCCGCTGGAAGAAAACAAAGAAGCTAACAAAGCTGAAGAATCTTCTTGCATCGAAAAATTCTCTCAGATCGTTCAGGAATGGCAGGGTAAAATCGCTGCTGTTATCATCGAACCGATCCAGTCTGAAGGTGGTGACAACCACGCTTCTTCTGACTTCTTCCAGAAACTGCGTGAAATCACCATCGAAAACGGTATCCTGATGATCGTTGACGAAGTTCAGACCGGTGTTGGTGCTACCGGTAAAATGTGGGCTCACGAACACTGGAACCTGTCTAACCCGCCGGACCTGGTTACCTTCTCTAAAAAATTCCAGGCTGCTGGTTTCTACTACCACGACCCGAAACTGCAACCGGACCAGCCGTTCCGTCAGTTCAACACCTGGTGCGGTGACCCGTCTAAAGCTCTGATCGCTAAAGTTATCTACGAAGAAATCGTTAAACACGACCTGGTTACCCGTACCGCTGAAGTTGGTAACTACCTGTTCAACCGTCTGGAAAAACTGTTCGAAGGTAAAAACTACATCCAGAACCTGCGTGGTAAAGGTCAGGGTACCTACATCGCTTTCGACTTCGGTACCTCTTCTGAACGTGACTCTTTCCTGTCTCGTCTGCGTTGCAACGGTGCTAACGTTGCTGGTTGCGGTGACTCTGCTGTTCGTCTGCGTCCGTCTCTGACCTTCGAAGAAAAACACGCTGACGTTCTGGTTTCTATCTTCGACAAAACCCTGCGTCAGCTGTACGGTTAA

#### BaPAT

ATGCATCATCACCATCACCACAACCAGCCGCTGAACGTTGCTCCGCCGGTTTCTTCTGAACTGAACCTGCGTGCTCACTGGATGCCGTTCTCTGCTAACCGTAACTTCCAGAAAGACCCGCGTATCATCGTTGCTGCTGAAGGTTCTTGGCTGACCGACGACAAAGGTCGTAAAGTTTACGACTCTCTGTCTGGTCTGTGGACCTGCGGTGCTGGTCACTCTCGTAAAGAAATCCAGGAAGCTGTTGCTCGTCAGCTGGGTACCCTGGACTACTCTCCGGGTTTCCAGTACGGTCACCCGCTGTCTTTCCAGCTGGCTGAAAAAATCGCTGGTCTGCTGCCGGGTGAACTGAACCACGTTTTCTTCACCGGTTCTGGTTCTGAATGCGCTGACACCTCTATCAAAATGGCTCGTGCTTACTGGCGTCTGAAAGGTCAGCCGCAGAAAACCAAACTGATCGGTCGTGCTCGTGGTTACCACGGTGTTAACGTTGCTGGTACCTCTCTGGGTGGTATCGGTGGTAACCGTAAAATGTTCGGTCAGCTGATGGACGTTGACCACCTGCCGCACACCCTGCAACCGGGTATGGCTTTCACCCGTGGTATGGCTCAGACCGGTGGTGTTGAACTGGCTAACGAACTGCTGAAACTGATCGAACTGCACGACGCTTCTAACATCGCTGCTGTTATCGTTGAACCGATGTCTGGTTCTGCTGGTGTTCTGGTTCCGCCGGTTGGTTACCTGCAACGTCTGCGTGAAATCTGCGACCAGCACAACATCCTGCTGATCTTCGACGAAGTTATCACCGCTTTCGGTCGTCTGGGTACCTACTCTGGTGCTGAATACTTCGGTGTTACCCCGGACCTGATGAACGTTGCTAAACAGGTTACCAACGGTGCTGTTCCGATGGGTGCTGTTATCGCTTCTTCTGAAATCTACGACACCTTCATGAACCAGGCTCTGCCGGAACACGCTGTTGAATTTTCTCACGGTTACACCTACTCTGCTCACCCGGTTGCTTGCGCTGCTGGTCTGGCTGCTCTGGACATCCTGGCTCGTGACAACCTGGTTCAGCAGTCTGCTGAACTGGCTCCGCACTTCGAAAAAGGTCTGCACGGTCTGCAAGGTGCTAAAAACGTTATCGACATCCGTAACTGCGGTCTGGCTGGTGCTATCCAGATCGCTCCGCGTGACGGTGACCCGACCGTTCGTCCGTTCGAAGCTGGTATGAAACTGTGGCAGCAGGGTTTCTACGTTCGTTTCGGTGGTGACACCCTGCAATTCGGTCCGACCTTCAACGCTCGTCCGGAAGAACTGGACCGTCTGTTCGACGCTGTTGGTGAAGCTCTGAACGGTATCGCTTAATCTAGAGCTAGCG

#### LeuDH^*B. cereus*^

ATGCATCATCACCATCACCACACCCTGGAAATCTTCGAATACCTGGAAAAATACGACTACGAACAGGTTGTTTTCTGCCAGGACAAAGAATCTGGTCTGAAAGCTATCATCGCTATCCACGACACCACCCTGGGTCCGGCTCTGGGTGGTACCCGTATGTGGACCTACGACTCTGAAGAAGCTGCTATCGAAGACGCTCTGCGTCTGGCTAAAGGTATGACCTACAAAAACGCTGCTGCTGGTCTGAACCTGGGTGGTGCTAAAACCGTTATCATCGGTGACCCGCGTAAAGACAAATCTGAAGCTATGTTCCGTGCTCTGGGTCGTTACATCCAGGGTCTGAACGGTCGTTACATCACCGCTGAAGACGTTGGTACCACCGTTGACGACATGGACATCATCCACGAAGAAACCGACTTCGTTACCGGTATCTCTCCGTCTTTCGGTTCTTCTGGTAACCCGTCTCCGGTTACCGCTTACGGTGTTTACCGTGGTATGAAAGCTGCTGCTAAAGAAGCTTTCGGTACCGACAACCTGGAAGGTAAAGTTATCGCTGTTCAGGGTGTTGGTAACGTTGCTTACCACCTGTGCAAACACCTGCACGCTGAAGGTGCTAAACTGATCGTTACCGACATCAACAAAGAAGCTGTTCAGCGTGCTGTTGAAGAATTTGGTGCTTCTGCTGTTGAACCGAACGAAATCTACGGTGTTGAATGCGACATCTACGCTCCGTGCGCTCTGGGTGCTACCGTTAACGACGAAACCATCCCGCAGCTGAAAGCTAAAGTTATCGCTGGTTCTGCTAACAACCAGCTGAAAGAAGACCGTCACGGTGACATCATCCACGAAATGGGTATCGTTTACGCTCCGGACTACGTTATCAACGCTGGTGGTGTTATCAACGTTGCTGACGAACTGTACGGTTACAACCGTGAACGTGCTCTGAAACGTGTTGAATCTATCTACGACACCATCGCTAAAGTTATCGAAATCTCTAAACGTGACGGTATCGCTACCTACGTTGCTGCTGACCGTCTGGCTGAAGAACGTATCGCTTCTCTGAAAAACTCTCGTTCTACCTACCTGCGTAACGGTCACGACATCATCTCTCGTCGTTAA

#### LeuDH^*L. sphaericus*^

ATGCATCATCACCATCACCACGAAATCTTCAAATACATGGAAAAATACGACTACGAACAGCTGGTTTTCTGCCAGGACGAAGCTTCTGGTCTGAAAGCTGTTATCGCTATCCACGACACCACCCTGGGTCCGGCTCTGGGTGGTGCTCGTATGTGGACCTACGCTTCTGAAGAAAACGCTGTTGAAGACGCTCTGCGTCTGGCTCGTGGTATGACCTACAAAAACGCTGCTGCTGGTCTGAACCTGGGTGGTGGTAAAACCGTTATCATCGGTGACCCGTTCAAAGACAAAAACGAAGAAATGTTCCGTGCTCTGGGTCGTTTCATCCAGGGTCTGAACGGTCGTTACATCACCGCTGAAGACGTTGGTACCACCGTTACCGACATGGACCTGATCCACGAAGAAACCGACTACGTTACCGGTATCTCTCCGGCTTTCGGTTCTTCTGGTAACCCGTCTCCGGTTACCGCTTACGGTGTTTACCGTGGTATGAAAGCTGCTGCTAAAGAAGCTTTCGGTTCTGAATCTCTGGAAGGTCTGAAAATCTCTGTTCAGGGTCTGGGTAACGTTGCTTACAAACTGTGCGAATACCTGCACAACGAAGGTGCTAAACTGGTTGTTACCGACATCAACCAGGCTGCTATCGACCGTGTTGTTAACGACTTCGACGCTATCGCTGTTGCTCCGGACGAAATCTACGCTCAGGAAGTTGACATCTTCTCTCCGTGCGCTCTGGGTGCTATCCTGAACGACGAAACCATCCCGCAGCTGAAAGCTAAAGTTATCGCTGGTTCTGCTAACAACCAGCTGAAAGACTCTCGTCACGGTGACTTCCTGCACGAACTGGGTATCGTTTACGCTCCGGACTACGTTATCAACGCTGGTGGTGTTATCAACGTTGCTGACGAACTGTACGGTTACAACCGTGAACGTGCTCTGAAACGTGTTGACGGTATCTACGACTCTATCGAAAAAATCTTCGCTATCTCTAAACGTGACGGTATCCCGACCTACGTTGCTGCTAACCGTCTGGCTGAAGAACGTATCGCTCGTGTTGCTAAATCTCGTTCTCAGTTCCTGAAAAACGAAAAAAACATCCTGCACGGTCGTTAA

#### BhcA

ATGCATCATCACCATCACCACACCTCTCAGAACCCGATCTTCATCCCGGGTCCGACCAACATCCCGGAAGAAATGCGTAAAGCTGTTGACATGCCGACCATCGACCACCGTTCTCCGGTTTTCGGTCGTATGCTGCACCCGGCTCTGGAAGGTGTTAAAAAAGTTCTGAAAACCACCCAGGCTCAGGTTTTCCTGTTCCCGTCTACCGGTACCGGTGGTTGGGAAACCGCTATCACCAACACCCTGTCTCCGGGTGACAAAGTTCTGGCTGCTCGTAACGGTATGTTCTCTCACCGTTGGATCGACATGTGCCAGCGTCACGGTCTGGACGTTACCTTCGTTGAAACCCCGTGGGGTGAAGGTGTTCCGGCTGACCGTTTCGAAGAAATCCTGACCGCTGACAAAGGTCACGAAATCCGTGTTGTTCTGGCTACCCACAACGAAACCGCTACCGGTGTTAAATCTGACATCGCTGCTGTTCGTCGTGCTCTGGACGCTGCTAAACACCCGGCTCTGCTGTTCGTTGACGGTGTTTCTTCTATCGGTTCTATGGACTTCCGTATGGACGAATGGGGTGTTGACATCGCTGTTACCGGTTCTCAGAAAGGTTTCATGCTGCCGCCGGGTCTGGCTATCGTTGGTTTCTCTCCGAAAGCTATGGAAGCTGTTGAAACCGCTCGTCTGCCGCGTACCTTCTTCGACATCCGTGACATGGCTACCGGTTACGCTCGTAACGGTTACCCGTACACCCCGCCGGTTGGTCTGATCAACGGTCTGAACGCTTCTTGCGAACGTATCCTGGCTGAAGGTCTGGAAAACGTTTTCGCTCGTCACCACCGTATCGCTTCTGGTGTTCGTGCTGCTGTTGACGCTTGGGGTCTGAAACTGTGCGCTGTTCGTCCGGAACTGTACTCTGACTCTGTTTCTGCTATCCGTGTTCCGGAAGGTTTCGACGCTAACCTGATCGTTTCTCACGCTCTGGAAACCTACGACATGGCTTTCGGTACCGGTCTGGGTCAGGTTGCTGGTAAAGTTTTCCGTATCGGTCACCTGGGTTCTCTGACCGACGCTATGGCTCTGTCTGGTATCGCTACCGCTGAAATGGTTATGGCTGACCTGGGTCTGCCGATCCAGCTGGGTTCTGGTGTTGCTGCTGCTCAGGAACACTACCGTCAGACCACCGCTGCTGCTCAGAAAAAAGC TGCTTAATCTAGAGCTAGC

